# Multi-omics approach reveals dysregulated genes during hESCs neuronal differentiation exposure to paracetamol

**DOI:** 10.1101/2022.12.08.519620

**Authors:** Mari Spildrejorde, Athina Samara, Ankush Sharma, Magnus Leithaug, Martin Falck, Stefania Modafferi, Arvind Y. M. Sundaram, Ganesh Acharya, Hedvig Nordeng, Ragnhild Eskeland, Kristina Gervin, Robert Lyle

## Abstract

Prenatal paracetamol exposure has been associated with neurodevelopmental outcomes in childhood. Pharmacoepigenetic studies show differences in cord blood DNA methylation between paracetamol exposed and unexposed neonates. However, causal implications and impact of long-term prenatal long-term paracetamol exposure on brain development remain unclear. Using a multi-omics approach, we investigated the effects of paracetamol on a model of early human brain development. We exposed human embryonic stem cells undergoing in vitro neuronal differentiation to daily media changes with paracetamol concentrations corresponding to maternal therapeutic doses. Single-cell RNA-seq and ATAC-seq integration identified paracetamol-induced chromatin-opening changes linked to gene expression. Differentially methylated and/or expressed genes were involved in signalling, neurotransmission, and cell fate-determination trajectories. Some genes involved in neuronal injury and development-specific pathways, such as *KCNE3*, overlapped with differentially methylated genes previously identified in cord blood associated with prenatal paracetamol exposure. Our data suggest that paracetamol may play a causal role in impaired neurodevelopment.

**Graphical Abstract:** 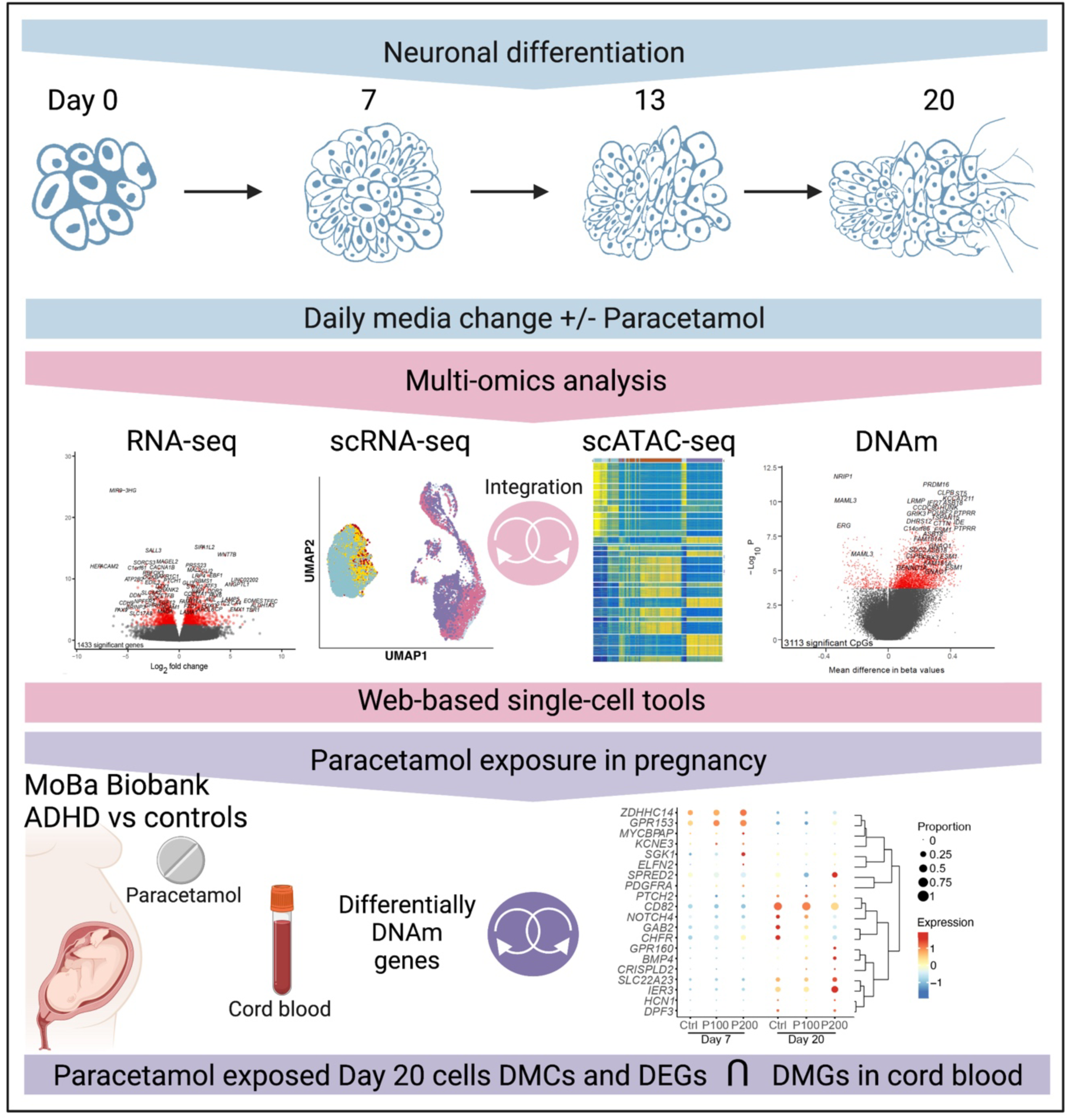

## Introduction

Paracetamol (also known as acetaminophen) is the most widely used analgesic and antipyretic during pregnancy, and it is considered safe for use as the first line option for pregnant women in need of mild analgesics or antipyretics^1–4^. A number of large epidemiological studies have reported an association between long-term maternal paracetamol use during pregnancy and increased risk of adverse neurodevelopmental outcomes, such as Attention Deficit/Hyperactivity Disorder (ADHD), in the child^5–13^. The association is reported to be stronger with long-term exposure and higher dose^14^. In 2019, the EUs pharmacovigilance safety committee (PRAC) reviewed all the available evidence, including non-clinical and epidemiological studies, regarding the impact of prenatal paracetamol exposure on impaired neurodevelopment in offspring. PRAC concluded that the available evidence is inconclusive, and recommended that the summary of product characteristics (SmPC) of paracetamol containing products should be updated to reflect the current state of scientific knowledge; “*Epidemiological studies on neurodevelopment in children exposed to paracetamol in utero show inconclusive results*” (PRAC, 2019)^15^. More recently, a group of researchers published a consensus statement and literature review in Nature Reviews Endocrinology concluding that there is growing evidence supporting the hypothesis that in utero exposure to paracetamol can impair fetal development^16^. This conclusion is still highly debated and contested by others (ENTIS, 2021)^17^,^18,19^ reflecting a need for further research.

Using samples from the Norwegian Mother, Father and Child cohort (MoBa) biobank, we have previously shown differences in DNA methylation (DNAm) in cord blood from children diagnosed with (ADHD) who have been exposed to paracetamol (>20 days) during prenatal development compared to unexposed children^20^. These findings suggest that DNAm might be involved in the pathogenesis of ADHD, but the causality and effect on neuronal differentiation and brain development is not known. It is well established that normal prenatal neurodevelopment involves cellular differentiation and establishment of cell-type specific epigenetic patterns, and that these events are prone to influences by environmental factors. For instance, maternal smoking has been shown to induce DNAm changes and modulate risk of neurodevelopmental disorders (NDDs)^21–23^. If and how paracetamol modulates the apparent increased risk of NDDs is currently unknown.

Recently, we established a protocol for neuronal differentiation of human embryonic stem cells (hESCs), which can be used in neuropharmacological studies^24^. In the present study, we have used this model system of early human brain development and investigated the effects of paracetamol exposures on transcriptional and epigenetic regulation. Paracetamol doses were selected to reflect therapeutic maternal doses and fetal in utero exposures^25–27^. By integrating multiple omics methods (bulk RNA-seq, bulk DNAm, single-cell RNA-seq and ATAC-seq) we observed time and dose effects of paracetamol exposure during neuronal differentiation.

## Results

### Neuronal differentiation exposure, timeline, and morphology

We investigated epigenetic and transcriptomic effects of exposure to paracetamol using an in vitro neuronal differentiation protocol that drives hESCs towards anterior neuroectoderm^24,28^. The neuronal differentiation is divided into three stages: the neural induction phase (Stage I) ends at Day 7, the self-patterning phase (Stage II) ends at Day 13, and the FGF2/EGF2-induced maturation phase (Stage III) ends at Day 20 (**Fig. 1A**). We replaced culture media daily, and the cells were exposed to 100 or 200 μM paracetamol during differentiation from Day 1 and onwards. These concentrations have been documented to be in the range of therapeutic plasma concentrations^25–27^. Unexposed (control) and paracetamol-exposed cells were harvested for downstream analyses on Day 7 and 20. We also harvested control cells at the onset of differentiation (hESCs; Day 0), and on the intermediate timepoint that cells were passaged (Day 13) to assess whether paracetamol exposed cells had mRNA abundance changes related to proliferation or delayed differentiation.

**Figure 1.**
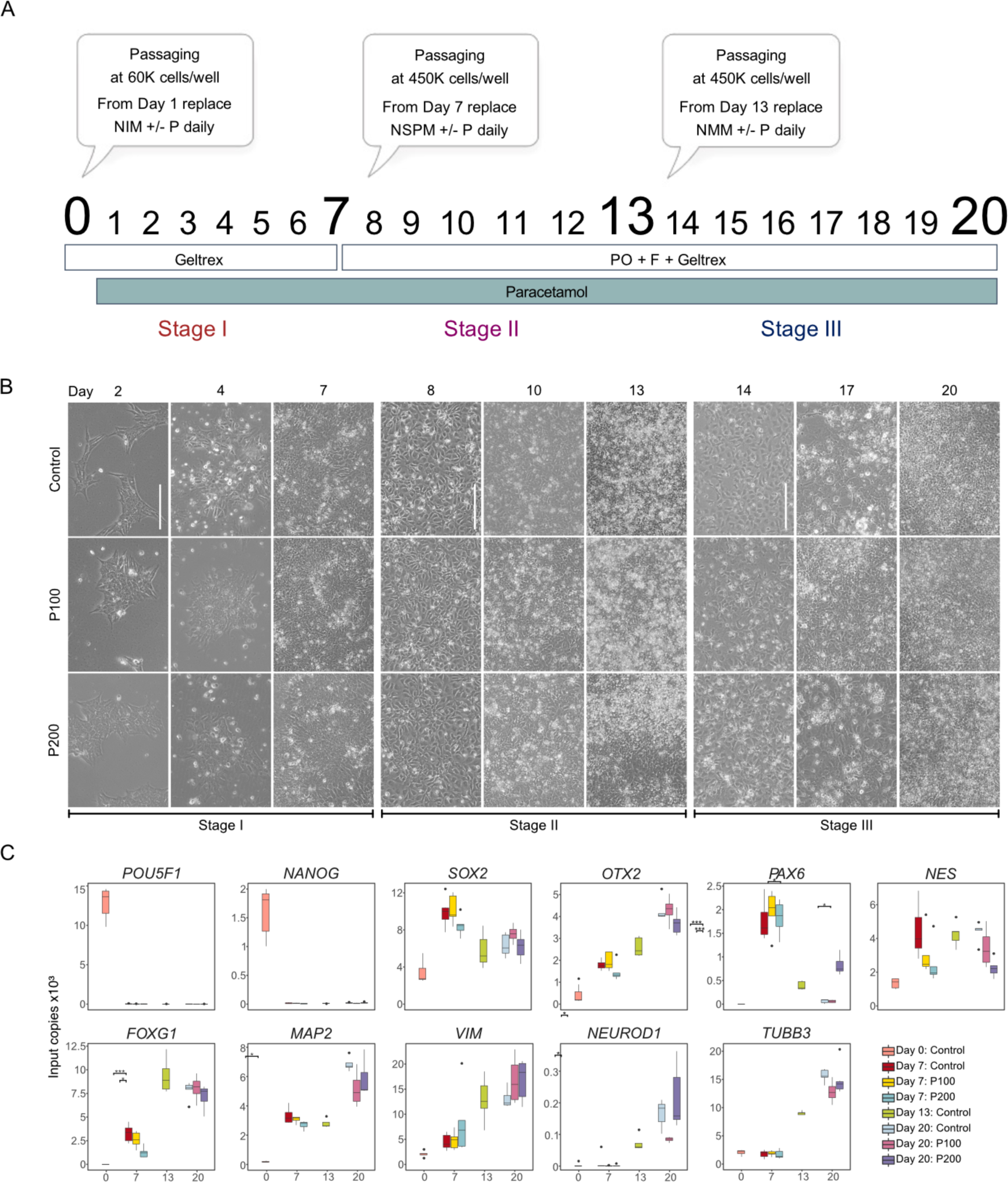
Neuronal differentiation of hESC to model neurodevelopmental effects of paracetamol. A) Schematic illustration of the different phases of the neuronal differentiation protocol from Day 0 to Day 20. Cells were exposed to 100 or 200 μM paracetamol from Day 1 and onwards. The effect of paracetamol on gene expression and epigenetic profiles was evaluated at Day 7 and Day 20. B) Representative brightfield images of the differentiation timeline for control, P100 or P200 cells during differentiation at Day 2, 4, 7, 8, 10, 13, 14, 17 and 20 (scale bar corresponds to 100 μM). C) ddPCR results from 4-6 replicates of mRNA expression of selected marker genes from Days 0, 7, 13 and 20. Significant comparisons are marked with asterisks (Student’s t-test, *: p <= 0.05, ***: p <= 0.001).

The timeline of brightfield images of the control cells versus cells exposed to 100 (P100) or 200 μM paracetamol (P200) documents the morphological changes and cell culture density at differentiation Days 2, 4, 7, 8, 10, 13, 14, 17 and 20 (**Fig. 1B**). Tightly packed neuroepithelial cells form the neural rosettes by Day 7 reassembled at the next stage under high density cell passaging and proceeded to maturation. We did not observe any distinct morphological changes in the differentiating cultures exposed to 100 (P100) or 200 μM paracetamol (P200). However, preliminary paracetamol titration experiments showed that exposing the differentiating cells to 400 μM paracetamol increased cell death and unpatterned morphology in cells and were thus discontinued. A set of representative images following the 20-day timeline in control cultures and 100, 200 and 400 μM paracetamol-exposed cells is presented in supplemental information (**Fig. S1A-C**).

### Validation of differentiation markers

The effect of paracetamol on gene expression was assessed at Days 7 and 20 was assayed using digital droplet PCR (ddPCR) (**Fig.1C**). Expression of the pluripotency transcription factors (TFs) *POU5F1* and *NANOG* decreased significantly after neural induction. As anticipated, we observed an increase in expression of the neural markers *SOX2, OTX2, FOXG1* and *MAP2* on Day 7 and *VIM, TUBB3* and *NEUROD1* on Day 13. Notably, the expression of *FOXG1, MAP2* and *NES* was significantly different between exposed and control cells on Day 7. Differential expression upon paracetamol exposure was also documented for filament *NES, PAX6* and *NEUROD1* on Day 20.

### Paracetamol-induced gene expression changes in genes involved in neural development

To delineate the effect on gene expression, we performed bulk gene expression analysis using RNA-seq in controls and paracetamol exposed cells (P100 and P200; **Fig. 2, S2** and **Table S1**). Principal component analysis showed that samples clustered according to differentiation day (**Fig. S2A; STAR Methods**). Overall, when we compared P100 and control we identified 121 differentially expressed genes (DEGs, **Fig. 2A**) and 1 433 DEGs between P200 and control (**Fig. 2B**) from Day 7 to Day 20 (**Table S1**). Pairwise comparisons between paracetamol (P100 and P200) and control at each day are shown in **Fig. S2B-C**. The bulk RNA-seq analysis of the previously selected marker genes (**Fig. 1B**) correlated well with the ddPCR results of selected marker genes (**Fig. S1D**).

**Figure 2.**
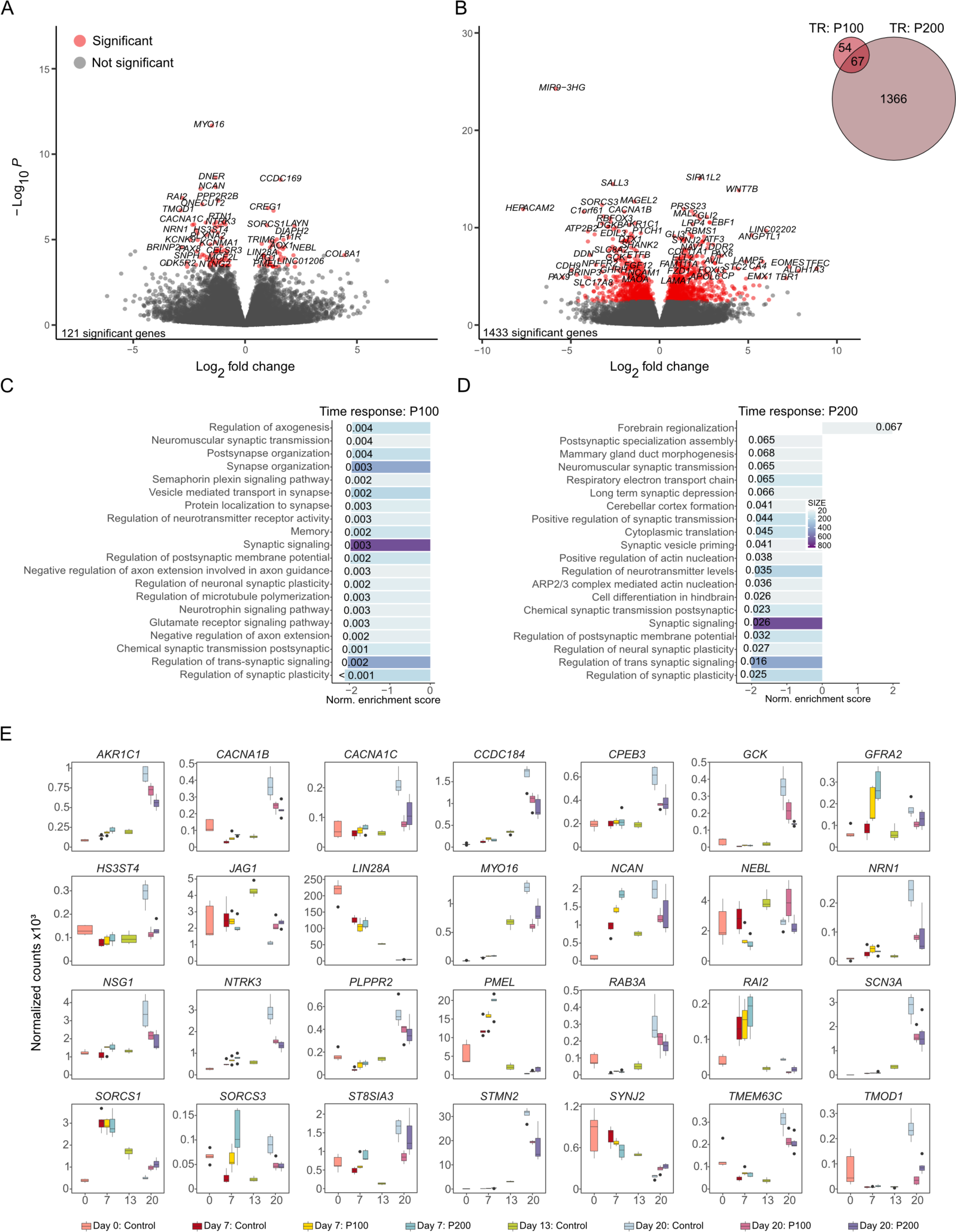
Paracetamol exposure of differentiating hESCs modulates expression of genes involved in neural development. Volcano plots of longitudinal differences in gene expression in P100 and B) P200 cells from Day 7 to Day 20. Corresponding Venn diagram shows the number of overlapping DEGs between the time-response (TR) of the two concentrations of paracetamol. Top 20 enriched BPs in C) P100 cells and D) P200 cells over time. E) Gene expression levels of selected DEGs, which overlapped between TR P100 and TR P200.

Gene set enrichment analyses (GSEA) analyses identified enrichment of downregulated BPs involved in synaptic organization, transmission, and regulation in the P100 time-response analysis (**Fig. 2C**). Notably, the upregulated BPs in the P200 time-response analysis were enriched for forebrain regionalization, cerebellar cortex formation and hindbrain differentiation (**Fig. 2D**). Also, the DEGs associated with P100 and P200 were enriched for common BPs reflecting enrichment of transmitter transport and regulation, synaptogenesis, synaptic organization, and plasticity.

The bulk RNA-seq pairwise comparisons of the P100 and P200 paracetamol-exposed cells to controls, showed an overlap of several DEGs (**Fig. S2D-F**). Exposure to paracetamol was associated with downregulation of genes previously linked to migration and neural development (e.g. *NTRK3, PMEL*^29^, *CCDC184*^30^, *MYT1*^31,32^). Furthermore, we identified DEGs linked to synaptic organization and transduction (e.g. *MYO16* ^33^, *HS3ST4* ^34^*, SORCS3* ^35^), metabolism (*GCK* ^36^), neuronal survival, dendrite branching, axonal growth and neural projection in development (e.g. *NSG1*^37^, *TMEM3C6*^38^, *TMOD1*^39^*, PLPPR4*^40^*, NEBL*^41^, *NRN1*^42^ and *GFRA2*^43^) and channelopathies (e.g. *CACNA1B/C*^44^, *SCN3A*^45^) (**Fig. 2E**). Thus, these bulk RNA-seq analyses revealed transcriptional dysregulation of genes related to possible developmental delays between control and paracetamol-exposed cells.

### Single-cell RNA sequencing reveals dose-specific changes in several major cellular processes after paracetamol exposure

To explore cell-type specific gene expression and maturation signatures over time, we performed single-cell RNA sequencing (scRNA-seq) analysis of control and paracetamol-exposed cells (P100 and P200) at Days 7 and 20 (**Table S2**). Specifically, our aim was to determine whether paracetamol exposure caused deviations in neuronal differentiation compared to control cells. A total of 15 201 cells (n=6 924 Day 7 cells, n=8 277 Day 20 cells) from two time-course experiments were aggregated and projected in Uniform Manifold Approximation and Projections (UMAPs, **Fig. 3; STAR Methods**). The scRNA-seq data may also be visualized in the open access webtool (**<u>hescneuroparacet</u>**), where expression of genes can be explored per cell, cluster, and time point.

**Figure 3.**
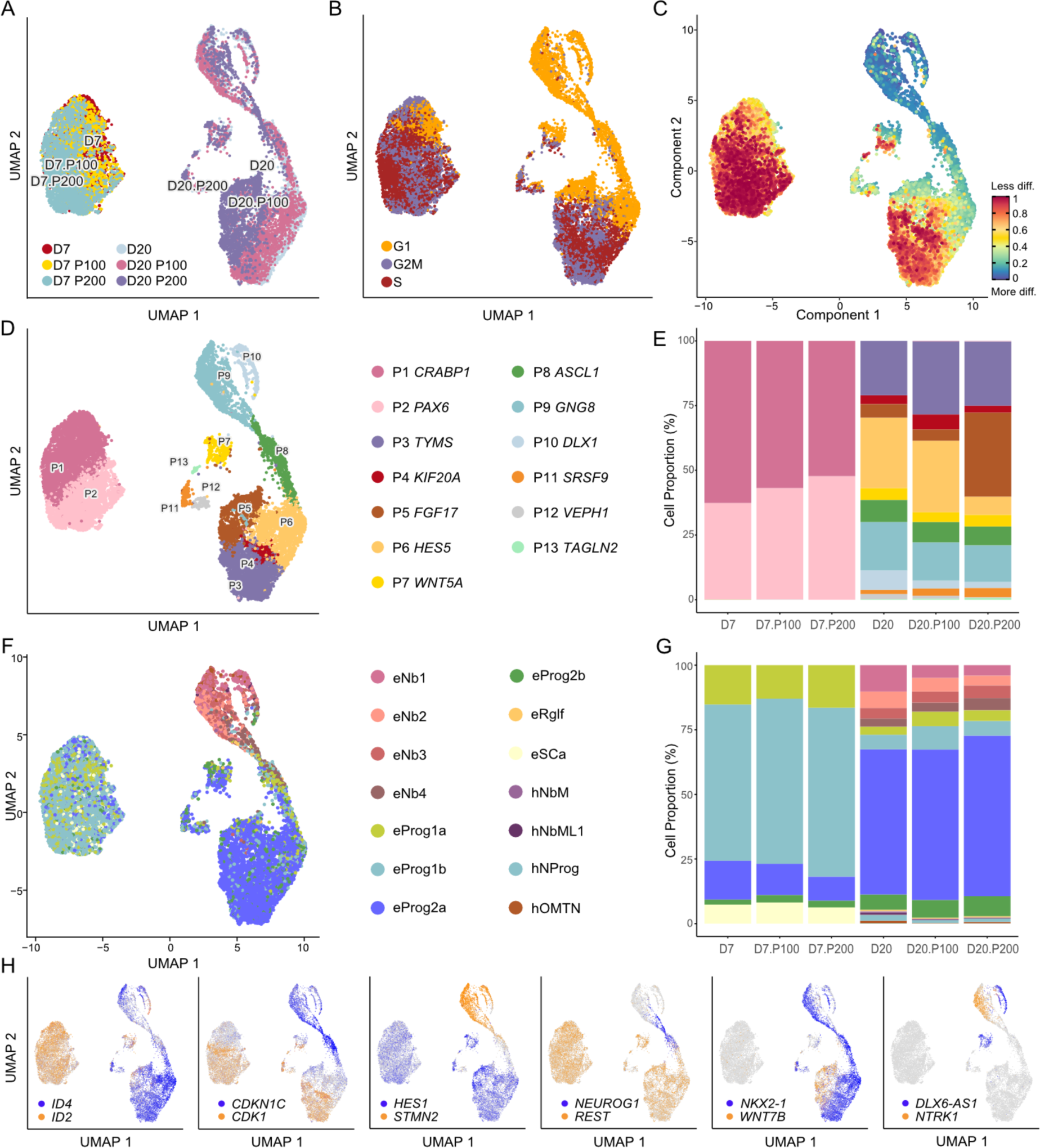
scRNA-seq analysis revealed shifts in cell type composition of differentiating cells exposed to paracetamol. Day 7 and Day 20 control cells and cells exposed to 100 μM (P100) or 200 μM (P200) paracetamol were visualized with UMAP and coloured by A) sample identity, B) Seurat predicted cell cycle phase, C) CytoTRACE pseudotime differentiation trajectory, D) defined Seurat clusters at resolution 0.4 with corresponding gene annotations and E) cell proportions per cluster. F) UMAP plots of cells coloured by SingleR cell annotation to Human Brain reference with corresponding cell annotations. Cell types starting with “e” are hESC derived cells and cell types starting with “h” are in vivo human embryo cell types. Nb1-4; neuroblasts, Prog1-2; neuronal progenitors, Rglf; radial glia-like cells, SCa; stem cells, NbM; medial neuroblasts, NbML1; mediolateral neuroblasts, NProg; neuronal progenitors, OMTN; oculomotor and trochlear nucleus. G) Corresponding cell proportions per cell type. H) UMAP plots of dual gene coexpression for *ID2/ID4, CDK1/CDKN1C, HES1/STMN2, REST/NEUROD1, NKX2.1/WNT7B* and *NTRK.1/DLX6-AS1* indicate that maturation signatures agree with the CytoTRACE trajectory analysis.

The cells clustered according to differentiation day and not exposure to paracetamol (**Fig. 3A**). Consistent with previous results^28^, seurat-predicted cell cycle phase showed an increase in G1 cells at Day 20 (**Fig. 3B**). However, the existing cell cycle analysis tools are unable to decipher the proportion of neurons that are in G0 phase. Using CytoTRACE pseudotime differentiation trajectory analysis, the most differentiated cells were found at Day 20 (**Fig. 3C**). The composite neuronal differentiation P1-P13 clusters were manually annotated with genes *CRABP1, PAX6, TYMS, KIF20A, FGF17, HES5, WNT5A, ASCL1, GNG8, DLX1, SRSF9, VEPH1* and *TAGLN2*, respectively (**Fig. 3D**). At Day 7, we observed a subtle shift in cell composition from P1 to P2 in the cells exposed to paracetamol (**Fig. 3E)**. The effect was more prominent at Day 20 with a higher proportion of paracetamol-exposed cells compared to controls annotated to cluster P5 and a more prominent effect at P200 exposure, whereas a lower proportion were annotated to P6, P9 and P10 for both concentrations (**Fig. 3E**).

To further investigate cell identity, the data were juxtaposed with a scRNA-seq Human Brain dataset^46^ (**STAR Methods**). Most cells at Day 7 were resembled neuronal progenitors, whereas cells at Day 20 were similar to neuronal progenitors, neuroblasts and neurons (**Fig. 3F**). At Day 20, there was a subtle shift from cells annotated as neuroblasts towards the less differentiated neuronal progenitors in paracetamol-exposed cells compared to controls (**Fig. 3G).** UMAP plots of the dual co-expression of the genes *ID2/ID4, CDK1/CDKN1C, HES1/STMN2, REST/NEUROD1, NKX2-1/WNT7B* and *NTRK1/DLX6-AS1* indicate agreement between these maturation signatures and the CytoTRACE trajectory analysis (**Fig. 3C** and **H**).

The analysis of the top DEGs per cluster (**Fig. S4**) and data exploration (**Fig. S5)**, showed that paracetamol exposure induced changes in several major processes at the selected timepoints. We identified dose-dependent changes that link paracetamol exposure to cell-cycle transition important in neuronal maturation (*MKI67, PCNA, TP53, CDK1, MYBL2* and *GJA1)* neural induction differentiation, or its inhibition (*REST, RAX, PAX6, HES1, HES5, ID3* and *ID4),* neurite outgrowth and cortical neurogenesis (*GBA2, ASPM*), neuronal maturation *(CDKN1C, POU2F1, POU3F1, ROBO1, STMN2* and *STMN4)* and WNT and FGF signalling (*FRZB, WNT4, WNT7B, FGF8* and *FGFRL1)* (**Fig. S5)**. We observed a differential and dose dependent expression of crucial spatiotemporally regulated transcription factors associated with brain development (e.g., *NKX2.1, OTX2, FOXG1, ASCL1, ISL1, EMX2* and *HOXA1*), neurotransmitter transporter expression (*SAT1, SLC2A1*). Finally, we found differential expression of genes related to cellular response to toxic insults (e.g., *DDR1*; **Fig. S5)**.

### Paracetamol exposure is associated with differential expression of neural lineage markers

To extract the biological significance and interpretation of DEGs after paracetamol exposure in the scRNA-seq datasets, we performed GO analyses and identified enrichment of GO terms and BPs. First, we identified the top 10 upregulated and downregulated BPs between control cells and P100 (**Fig. S3A**) or P200 cells (**Fig. S3B**) on Day 7. Notably, BPs involved in *DNA replication* and *cell-cycle regulation* were upregulated in cells exposed to both paracetamol doses, and a downregulation of a BP which involves *generation of neurons*. DEGs in P100 cells compared to control cells were enriched for more neuronal specific annotations. Furthermore, P200 compared to control cells identified upregulated GO terms involved in cellular responses to toxic insults and *DNA checkpoint activation*. The top 20 DEGs between P100 (**Fig. S3C**) or P200 (**Fig. S3D**) and control cells at Day 7, are shown as bubble plots of relative gene expression. The gene expression of the top overlapping genes between the P100 and P200 cells compared to control cells at Day 7 (**Fig. S3E**) extends the GO annotations to gene specific similarities. We identified changes in crucial genes, such as the master gene of forebrain development *FOXG1*^47^ and genes of the *HES* and *ID* gene families involved in differentiation and neurogenesis^48^.

Next, we extracted the top 10 upregulated and downregulated BPs among the DEGs between control cells and P100 (**Fig. S3F**) or P200 cells (**Fig. S3G**) on Day 20. In both comparisons, downregulated GO terms included *neuron/nervous system developmen*t and *microtubule polymerization or depolymerization*. Of the top 20 DEGs at Day 20 between paracetamol-exposed and control cells, we found major patterning TFs, such as *NKX2-1* and *EMX2* (**Fig. S3H-I)**. Moreover, genes that belong to the ZIC family, among other genes involved in Notch and Wnt signalling, were also identified (**Fig. S3H-I)**. The gene expression of the top overlapping genes between P100 and P200 cells were compared to control cells at Day 20 (**Fig. S3J**). The analysis of GO terms delineated gene specific changes, such as upregulation of *SELENOW* (**Fig. S3D**), previously associated with neuroprotection from oxidative stress^49^. The paracetamol-induced differentiation lag as evidenced by the *PAX6* expression in the P200 cells (**Fig. 1C**). We also observed a downregulation of tissue- and stage-specific genes, such as *FABP7, ISL1, STMN2* and *INA*. In addition, *TUBB1A*, the isotype associated with postmitotic neurons^50^, and *TUBB2B*, that constitutes 30% of all brain beta tubulins^51^, were also found among the downregulated genes. Interestingly, *PEG10* and *C1orf61* and *MIR9-1HG* appear in the Day 20 P200 comparisons, genes which have recently been linked to cortical migration and intercellular communication^52^, further documenting how the P200 dose of paracetamol could affect proper network formation and cell-to-cell signalling.

### Integration of scATAC- and scRNA-seq link paracetamol-induced changes in chromatin opening to transcriptional activity at Day 20

To understand whether paracetamol exposure during differentiation influenced the chromatin landscapes at Day 20 we performed scATAC-seq. We obtained 3 042 nuclei for Day 20 control and 3 480 and 4 282 nuclei for Day 20 exposed to 100 μM (P100) and 200 μM paracetamol (P200), respectively. First, we reanalysed scRNA-seq data from Day 20 controls, P100 and P200, and remapped the P3-P13 clusters (**Fig. 4A-B**). The maturation trajectory cohered with the initial Day 7-Day 20 time point analysis (**Fig. 4C and 3C**). Next, we mapped the scATAC-seq data (**Fig. 4D**) to 15 scATAC-seq clusters (C1-C15; **Fig. S6A**) that we integrated with the annotated scRNA-seq P3, P5, P9, P10 and P13 clusters (**Fig. 4E)**. The quality of the combined scATAC-seq datasets was documented with an even distribution of integrated clusters (P3, P5, P9 and P10) over TSS, promoters, exons, introns, and distal genomic regions (**Fig. S6B-C**). The P13 cluster is represented by very few nuclei and displayed lower enrichment across all genomic regions.

**Figure 4.**
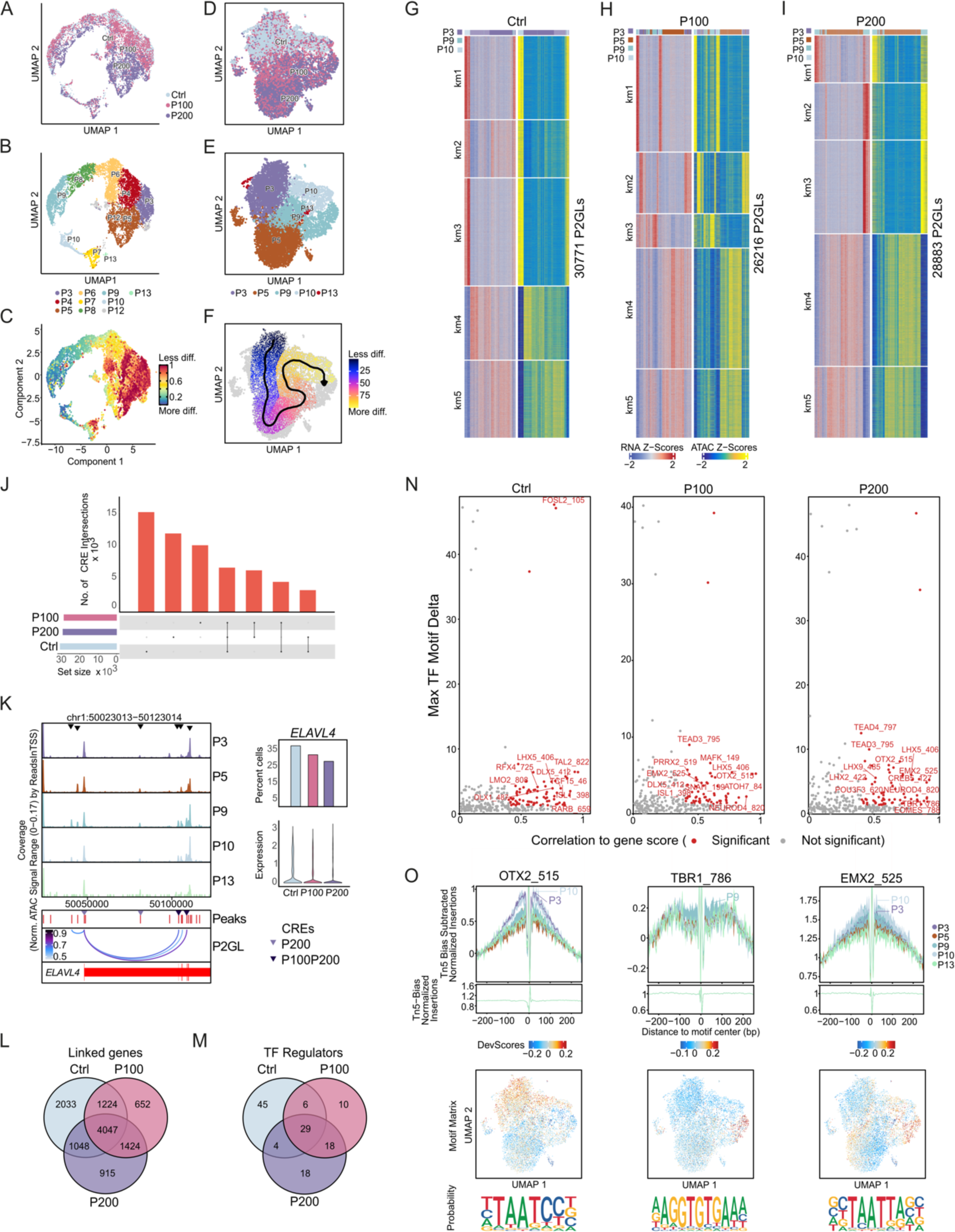
Effects of paracetamol on chromatin accessibility and integration with scRNA-seq at Day 20. A-C) scRNA-seq UMAP plots coloured by A) sample identity, B) scRNA-seq clusters or C) CytoTRACE pseudotime differentiation trajectory. D-E) scATAC-seq UMAP plots of coloured by D) sample identity and E) remapped clusters following constrained alignment of cell populations by scATAC-seq and scRNA-seq integration. F) Supervised pseudotime trajectory of integrated clusters. G-I) Heatmaps of scATAC-seq and scRNA-seq side-by-side representing peak to gene links (P2GLs) in G) control cells, H) P100 cells and I) P200 cells. Rows were clustered using k-means (k = 5). J) Overlap of putative CREs from P2GLs analyses of Day 20 control versus P100 and P200. K) *ELAVL4* locus browser view (GRCh38.p13) from 5000 cells with ATAC-seq signals in integrated clusters. Differential chromatin opening (black triangles), ATAC peaks (red), P2GLs (arcs) and putative CREs enriched in P200 or P100 and P200 cells (coloured triangles) are shown. The bar plot to the right shows the corresponding percentage of cells expressing *ELAVL4*, and the plot below represent the expression levels. L) A Venn diagram representing overlap of linked genes. M) Venn diagram of TF regulators identified from P2GLs integrative analysis. N) A selection of positive TF regulators computed using gene integration scores with motifs in the putative CREs. O) Footprints for selected TF regulators OTX2, TBR1 and EMX2 demonstrating preferential opening per cluster; below are the corresponding ArchR motif deviation scores, motif matrix and representative sequence logos identified in the scATAC-seq dataset.

To further explore the differential chromatin accessibility across the genome, we correlated distal accessible regions with gene expression^53^. This allows for identification of putative cis regulatory elements (CREs) for biological and functional comparisons of Day 20 control cells with P100 and P200 exposed cells. We grouped the peak-to-gene links (P2GLs) into five clusters and plotted heatmaps of gene expression and gene scores for all three datasets (P2GLs represent potential genes linked to chromatin opening, **Fig. 4G-I**). We observed relatively similar numbers of putative CREs in controls (n=30 771), P100 (n=26 216) and P200 (n=28 883). Notably, the heatmaps of controls and P100 and P200 cells were different with an absence or presence of the integrated cluster P5, also described as having differential prominence in the scRNA-seq data (**Fig. 3E**). Only 6 960 of the putative CREs overlapped between the three datasets, whereas 4 732, 3 435 and 6 534 putative CREs specifically overlapped between control and P100, control and P200, and P100 and P200, respectively (**Fig. 4J** and **Table S3A-C**). Moreover, we observed that many of the putative CREs were only detected in individual datasets (Day 20 control; n=15 644, P100; n=10 460 and P200; n=12 304 CREs). These variations in putative CREs suggests that paracetamol exposure results in changes in chromatin accessibility. For example, in the locus of the neuronal marker gene *STMN2,* which showed higher gene expression in unexposed cells compared to exposed cells, showed changes in chromatin opening peaks in paracetamol exposed cells (**Fig. S6D, S3J** and **S5**). Furthermore, accessibility peaks in the *ELAVL4* locus displayed differential chromatin opening in clusters P3, P5, P9 and P10. We also identified putative CREs in the exposed cells that correlated with higher *ELAVL4* expression in control cells compared to P100 or P200 cells (**Fig. 4K** and **Table S3A-C)**.

### Paracetamol affects region-specific chromatin accessibility

The putative enhancer-gene interactions identified likely represent chromatin regulomes in the control and the paracetamol-exposed cells. We therefore compared the level of overlap of linked genes to better understand the effect of paracetamol exposure on chromatin regulation. A larger proportion of linked genes overlapped between the Day 20 control and paracetamol exposed cells (n=4 047), and the P100 (n=1 224) and P200 (n=1 048) cells (**Fig. 4L** and **Table S3A-C** and **V**). GO analyses of the linked genes revealed a common enrichment of BPs, such as *nervous system development, biological regulation, neurogenesis,* and *neuron differentiation*, varying in the different k-means (km) clusters (**Fig. S6D** and **Table S3D-F**). Linked genes identified in paracetamol exposed cells (P100, n = 652; P200, n = 915 and 1424 for both P100 and P200) indicate exposure-induced changes in gene expression. Interestingly, linked genes including neuronal lineage transcription factors (e.g. *PAX6, NEUROD4, NEUROG1, SOX9* and *SOX2*) and genes involved in chromatin modification (e.g. histone H3K27 acetyltransferase *EP300*^54^, Histone H3K27me3 demethylase *KDM6A*^55^ and chromatin remodeller SNF2 subfamily member *SMARCAD1*^56^), suggest that paracetamol may contribute to modulation of transcriptional regulation and chromatin structure (**Table S3V**). In addition, GO analysis of the putative CREs identified enrichment of BPs per k-means group and sample are shown in **Tables S3G-U**.

Enriched TF motifs in the CREs may infer gene expression program regulation. We identified TFs (n=29) that were common for control, P100 and P200 cells, including *NEUROD1, NEUROG1, SOX9, HMGA1* and members of the ONECUT and NHLH families, which have previously been described in the neuronal differentiation protocol^28^. TFs that were common between P100 and P200 cells (n=18) or dependent on paracetamol dose (**Fig. 4M-N, S3I, S5, S6F** and **Table S4**). The TF footprint per cluster and motif enrichment and computed sequence logos for *OTX2, TBR1* and *EMX2* show that these factors are enriched and potentially regulate genes in these integrated clusters (**Fig. 4O)**.

### Paracetamol induced changes in DNAm during neuronal differentiation

To assess whether exposure of hESCs undergoing neuronal differentiation to paracetamol induced DNAm changes, we analysed control cells and cells exposed to 100 (P100) and 200 μM paracetamol (P200) at Day 7 and 20. The experimental set-up included analysis of control cells harvested at Day 0 and 13 as reference points of possible dysregulation (**Fig. 5** and **S7, Table S1**). Assessment of principal component analysis showed clustering of samples according to differentiation day (**Fig. S7C; STAR Methods**). Overall, the distribution of DNAm was indistinguishable across exposed and control cells at all time points regardless of paracetamol dose (**Fig. S7A**). In contrast, non-CpG DNAm levels decreased during differentiation and were lower in cells exposed to 200 μM paracetamol compared to control cells at Day 20 (**Fig. S7B**).

**Figure 5.**
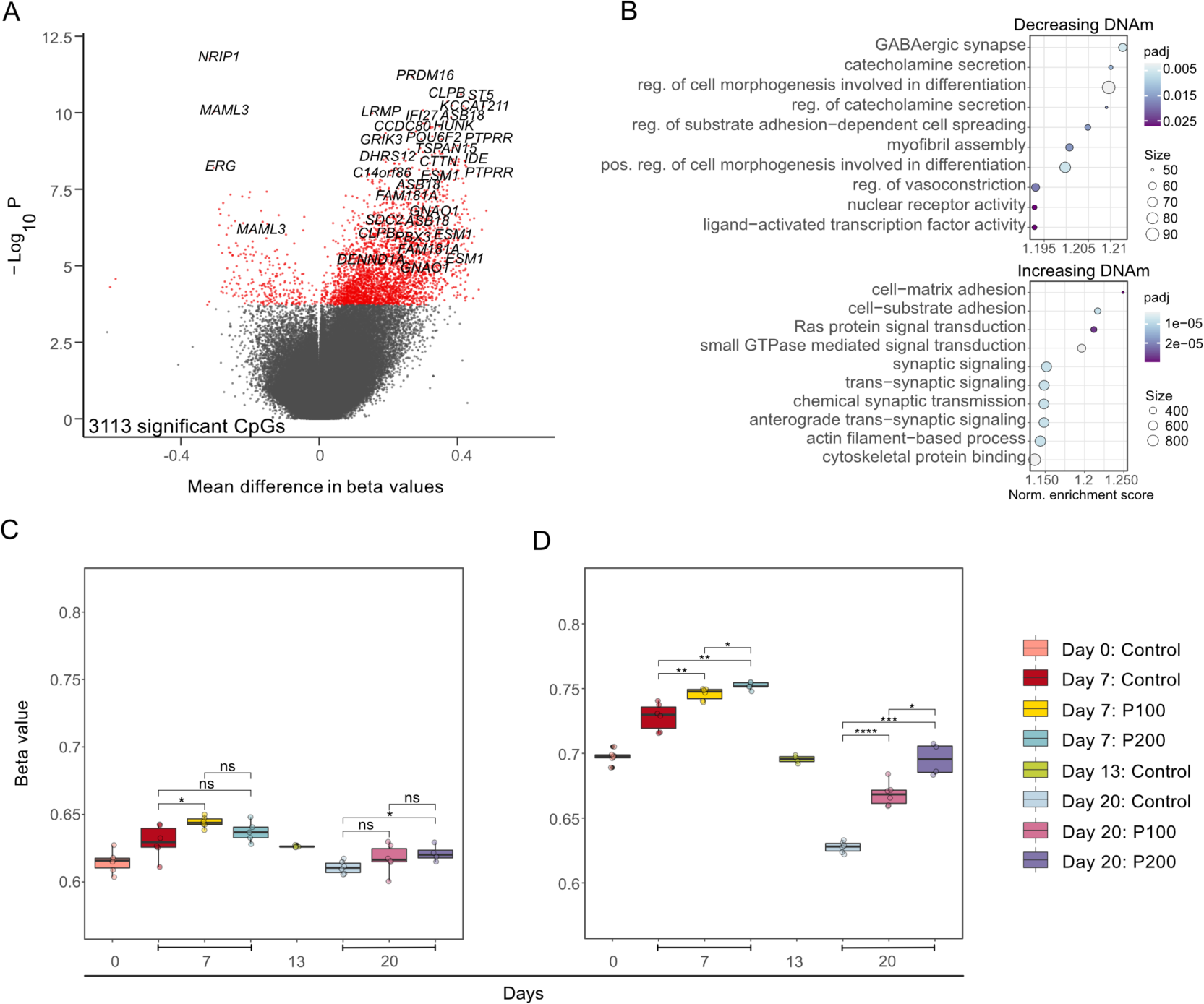
Exposure to paracetamol induces changes in DNAm over time during neuronal differentiation. A) Volcano plot showing the effect of 200 μM paracetamol from Day 7 to Day 20. CpGs with adjusted p-value < 0.05 were considered significant. B) Corresponding enriched GO-terms for CpGs showing a decrease in DNAm (top) or an increase in DNAm (bottom) over time (Day 20 – Day 7) in P200 cells compared to control cells. C) Average DNAm levels per sample for all CpGs across differentiation for Day 0, 7, 13, and 20. D) Average DNAm levels per sample for all significant CpGs (paracetamol-exposed cells vs control comparisons) at Day 0, 7, 13, and 20. E-F) *p < 0.05, **p < 0.01, ***p < 0.001, ****p < 0.0001.

To investigate whether the cells exposed to different doses of paracetamol (P100 and P200) respond differently than control cells from Day 7 to Day 20, we performed a DNAm time-response analysis. We observed no significant DNAm changes in P100 compared to control between Day 7 and Day 20 (not shown). In contrast, 3 113 CpGs responded differently to P200 compared to control cells over time (**Fig. 5A**). CpGs showing an increase in DNAm are annotated to genes with key functions in dynamic cellular redox changes in the developing brain, such as the neural specification gene *PRDM16*^57^ enriched for GO terms such as synaptic signalling and chemical synaptic transmission (**Fig. 5B**). CpGs showing a decrease in DNAm are annotated to genes enriched for gene ontology (GO) terms involved in synaptic regulation, GABAergic signalling, and cell morphogenesis (**Fig. 5B**). Overall, when we assessed DNAm levels at all CpGs (**Fig. 5C**) compared to all significant CpGs (**Fig. 5D**), we found a general dose-dependent increase in DNAm levels at significant sites in paracetamol-exposed cells compared to controls at both Day 7 and Day 20. The annotation of differentially methylated CpGs (DMCs) in relation to genes and CpG islands was similar across the different comparisons (**Fig. S7D-E**).

Next, we performed comparisons of DNAm levels in paracetamol-exposed cells to controls at Day 7 (**Fig. S7F and H, Table S1**) and Day 20 (**Fig. S3G and I, Table S1**) for the different doses. As expected, we observed more significant DNAm changes after longer exposure (Day 20) and at higher concentration of paracetamol (P200) (**Table S1**). We found no dose-dependent DMCs at Day 7 whereas at Day 20 a larger number of CpGs (n=8 940) were differentially methylated between P100 and P200. Moreover, there was some overlap between DMCs at 7 and Day 20 (**Fig. S7J**). To define a regulatory role of paracetamol induced DNAm changes on gene expression, we assessed the overlap between DEGs and differentially methylated genes (DMGs) for the pairwise comparisons and time response analyses (**Table S1**). The percentage of DMGs overlapping with DEGs varied between 3 and 63 % for the different comparisons. The P200 time response analysis revealed 180 overlapping genes, and some selected genes are visualized in **Figure 6**. DNAm levels for *GRIK3, CACNA1D, ABAT, MAPT* and *ANKRD6* were inversely correlated with gene expression, whereas DNAm levels for *PAX7, CDH2* and *WNT7B* were positively correlated with gene expression at Day 20. *GLI3* had DNAm levels that correlated with both negative and positive regulation (**Fig. 6**).

**Figure 6.**
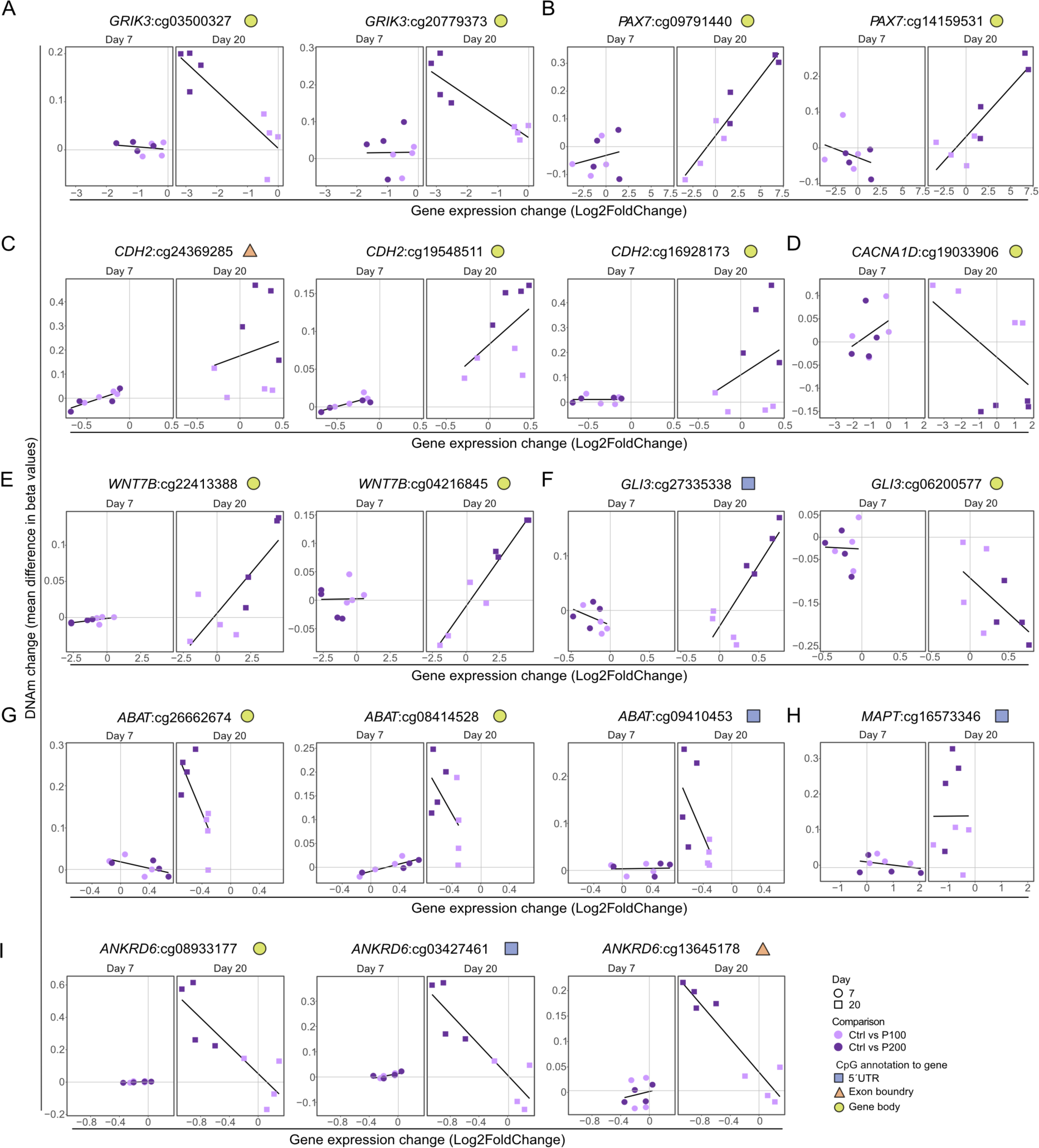
DNAm and gene expression for selected overlapping DMCs and DEGs. Change in DNAm and gene expression levels between control and P100 or P200 for A) *GRIK3*, B) *PAX7*, C) *CDH2*, D) *CACNA1D*, E) *WNT7B*, F) *GLI3*, G) *ABAT*, H) *MAPT* and I) *ANKRD6*. Each point represents a matched replicate between RNA-seq and DNAm and black lines represents a linear regression line of the mean of values.

### Overlap of dysregulated genes in differentiating hESCs and cord blood from children exposed to paracetamol during pregnancy

We have previously identified an association between differential DNAm in cord blood and long-term paracetamol exposure during pregnancy in children with ADHD^20^. To assess the translational potential and causality of these findings, we compared the dysregulated genes identified in the present model to the differentially methylated genes associated with paracetamol exposure in cord blood. In brief, the DMCs and the DEGs in paracetamol-exposed differentiating cells at Day 20 were correlated with DMGs in cord blood^20^. Interestingly, we identified an overlap of 20 genes between DEGs and DMCs for P100 and P200 in differentiating hESCs and differentially methylated genes identified in cord blood between paracetamol-exposed children with ADHD versus controls (**Fig. 7A**) and only one gene (*KCNE3*) with paracetamol-exposed children with ADHD versus ADHD controls (**Fig. 7B**).

**Figure 7.**
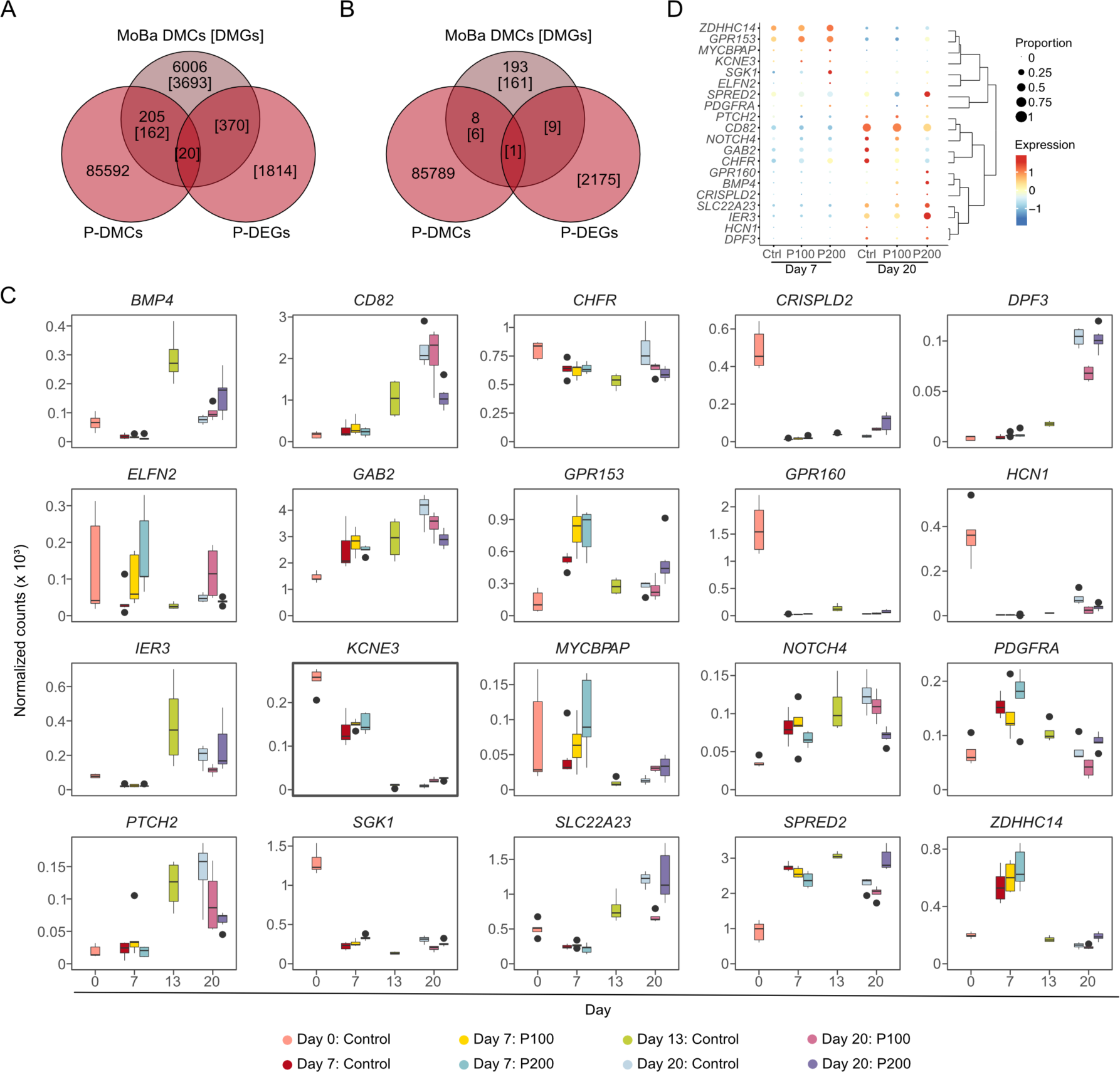
Overlap with differentially methylated genes in cord blood. A-B) Venn diagrams showing the overlap between DEGs and DMCs for P100 and P200 and A) paracetamol-exposed children with ADHD versus controls and B) paracetamol-exposed children with ADHD versus ADHD controls in cord blood. C-D) Gene expression levels of overlapping genes identified in cord blood derived from C) RNA-seq and D) scRNA-seq data.

We assessed the expression of these 20 overlapping genes in scRNA-seq data (**Fig. 7C**) and RNA-seq data (**Fig. 7D**). Notably, genes involved in differentiation, such as *GAB2*^58^, or *Notch and Hedgehog signalling pathways* (for example, *NOTCH4*^59^, *PTCH2*^60^ *SHISA2*^61^) show a dose-dependent downregulation of expression in paracetamol-exposed cells compared to controls. This high variation in expression levels after paracetamol exposure compared to controls, indicates a dose effect. Among genes identified in Day 7 P200 cells, we observed upregulation of *ZDHHC14*^62^ and *SGK1*^63^, that are associated with control of neuronal excitability and neuronal response to injury, respectively. Several toxic insult response genes were also upregulated at Day 20 in the P200 treated cells, such as *IER3*^64^, *SPRED2*^65^, *GPR130*^66^, and *SLC22A23*^67^.

## Discussion

To our knowledge, this is the first multi-omics study investigating how paracetamol exposure affects epigenetic and transcriptional programmes involved in early brain development using an in vitro hESC model of neuronal differentiation^28^.

We identified differential expression of genes enriched for BPs involved in transmitter transport and regulation, synaptic organization and synaptogenesis and plasticity. Interestingly, in cells exposed to the high dose of paracetamol (P200), we also identified enrichment of GO categories reflecting possible patterning deviations from forebrain differentiation. Thus, we observe an effect of paracetamol exposure on transcriptional dysregulation and possible developmental delays during neuronal differentiation. Additionally, among the common DEGs, irrespective of paracetamol dose, we identified *SORCS3,* which encodes a brain-expressed transmembrane receptor associated with neuronal development and plasticity that has been previously identified in a GWAS meta-analysis of ADHD significant risk loci^68^. The scRNA-seq analysis identified dose-dependent DEGs involved in several major processes during neuronal differentiation. Exposure to paracetamol resulted in a subtle shift in cell populations from P1 to P2 at Day 7. At Day 20 the P6, P9 and P10 clusters decreased whereas P5 markedly increased in P200. The DEGs link paracetamol exposure to genes involved in cell-cycle length and phase transition and genes important for neuronal maturation, neurite outgrowth, cortical neurogenesis, expression of neurotransmitter transporters, neuronal maturation and WNT and FGF signalling. Further, the data also documented dose-dependent differential expression of crucial spatiotemporally regulated TFs associated with brain development. These findings provide evidence of transcriptomic dose-dependent effects of paracetamol exposure related to cellular response to toxic insults and fate-determination deviation queues at the Day 20. Furthermore, the results identified an impact of paracetamol exposure on the regulation of central, autonomic and sympathetic nervous system development, which are all central to finetuning human cognition and associated with ADHD^69–71^.

To further characterize and investigate epigenetic mechanisms involved in altered cellular heterogeneity and differentiation, we performed scATAC-seq to map CREs and compare the chromatin landscape changes between control and cells exposed to P100 and P200 at Day 20 (**<u>hescneurodiffparacet</u>**). Interestingly, the results from these analyses identified differences in several chromatin accessible regions in genes previously associated with fate mapping and neurogenesis, which have been identified in risk determination studies focusing on cognition and autism spectrum disorders (e.g. *DLK1, DIO3, IPW* ^72,73^) and ADHD (e.g. *MEG3*^74^).

Genome-wide association studies (GWAS) suggest that cognitive disorders might also result from the cumulative effect of gene variant regulation and parent-of-origin effect^74^, which in ADHD has been associated with genes such as *DDC, MAOA, PDLIM1* and *TTR*. We also identified chromatin openness differences related to genes identified in the paracetamol exposed cells. For example, *PCDH7*, which acts as a cue for axonal guidance via roles in cell adhesion, was among the genes that were both differentially expressed and had different chromatin openness in Day 20 P200 cells. Dysregulation of *PCDH7* could be relevant to the semaphorin-plexin signalling documented by the enriched BPs in the P100 cells, and it is noteworthy as it has also been associated to ADHD^68^.

We also identified dose-dependent differences in chromatin accessibility of the neural specification gene locus *PRDM16*, which correlates well with DNAm analyses where exposure to paracetamol was associated with increased DNAm at CpGs annotated to *PRDM16*. This gene has been linked to regulation of transcriptional enhancers activating genes involved in intermediate progenitor cell production and repressing genes involved in cell migration^57^. The molecular mechanism of *PRDM16* remains unknown but has been associated with reactive oxygen species regulation in the embryonic cortex^75^.

Integrative analyses of scRNA-seq and scATAC-seq to explore the links between chromatin accessibility and gene expression documented high correlation and showed an overlap between scATAC-seq and scRNA-seq cell clusters. This analysis identified novel putative CREs, and linked genes affected by paracetamol exposure. Although a portion of linked genes overlapped between the control and exposed cells, there was only a small overlap between the putative CREs. This suggest that many local changes in chromatin opening were affected by paracetamol exposure and that these CREs may regulate different gene expression programmes in control and paracetamol exposed cells. How paracetamol regulates chromatin opening is not known, and we have limited knowledge on the effect of paracetamol on the TFs recognition of DNA motifs. However, our data suggest that genes linked to putative CREs that were affected by paracetamol exposure have a role in regulation of transcription and chromatin. In particular, a linked gene specific for paracetamol exposure was *EP300*, which codes for a histone acetyltransferase responsible for the active enhancer mark H3K27ac^76^. *EP300* expression decreased subtly upon paracetamol exposure both at Day 7 and 20, and these changes in EP300 levels may change the number of active enhancers in the paracetamol exposed cells. Furthermore, we identified TFs such as EMX2, TBR1 and OTX2 whose regulatory activity is affected by paracetamol exposure at the differentiation end point. This is suggestive of a mechanism for how paracetamol exposure may change gene expression programmes in early brain development. More work is needed to follow up the functional role of paracetamol exposure on the enhancer regulation and gene expression.

We found a general dose-dependent increase in DNAm in paracetamol exposed cells compared to controls at both Day 7 and Day 20. The genes were enriched for BPs involved in cell morphogenesis and adhesion, but also synaptic transmission and signalling with a focus on catecholamines and GABAergic transmission. Considering that this is an in vitro system, the findings are compelling since structural cortical changes and catecholaminergic transmission have been the target of various epidemiological studies and drug clinical trials, both in children and adults with ADHD^77–80^. It is biologically plausible that medications that cross the placenta and the blood-brain barrier, may interfere with normal fetal brain neurodevelopment, previously shown for valproic acid, and more recently for topiramate^81^. This is especially relevant for substances that cross the blood-brain barrier, which is considered functional by the 8th week of gestation^82^. However, invasive testing of the fetal compartment in human is rarely indicated, which makes data on safety during pregnancy for most substances scant. Thus, most pharmacoepigenetic studies address associations of DNAm differences in cord blood or placenta from neonates delivered at term exposed to the maternal medication during pregnancy, which is also the case for paracetamol^83–85^.

We previously found differentially methylated genes in cord blood from children with ADHD exposed to long-term maternal paracetamol use^20^. However, the causality, and relevance of such findings to brain development are not known. The hESCs used in our experiments represent the inner cell mass (ICM) of Day 6 preimplantation-blastocysts^86^. The ICM is known to differentiate to form the primitive ectoderm around Day 7.5 post fertilization^87^, but placental circulation is not established before Day 17-20^88^. Prior to this, the fraction of paracetamol that reaches the developing embryo does so via passive diffusion^89^. Interestingly, irrespective of the temporal differences of the two models^90^, comparing the differentially expressed and methylated genes identified in our study revealed overlap of several genes identified in cord blood with potential compelling relevance to early brain development. Notably, identification of a dose-specific effect on several genes have previously been shown to be associated with neural injury and toxic-insult response^91–93^. To our knowledge this is the first study that has identified an effect of paracetamol on differential DNAm of *KCNE3* in a neuronal differentiation model of hESCs. Differential DNAm at *KCNE3* was also identified in our MoBa study in cord blood^94^. Furthermore, *KCNE3* was also identified in a DNAm analysis of extremely low gestational age new born (ELGAN) cohort^83^. *KCNE3* is an interesting candidate, which encodes a voltage-gated ion channel with important functions in regulating release of neurotransmitters and neuronal excitability^95,96^. Among the overlapping genes, there was also *IER3*, whose differential DNA methylation at specific CpG sites has been previously associated with in utero exposure to bisphenol A (BPA)^97^. The same study had identified *TSPAN15*, which is also significantly differentially methylated between control and P200 cells in our neuronal differentiation model.

We also compared our data to previously published studies using the webtool (**<u>hescneuroparacet</u>**). In a rat model of fetal paracetamol exposure^98^, dysregulation in specific ABC efflux transporters and related enzymes was found after chronic treatment of the mothers in E19 fetal brain and choroid plexus. In our model of paracetamol exposure, we also identified differential expression of the relevant genes *ABCA1, ABCC5, ABCD3* and *GSTM*3. Moreover, the expression of *ABCC4*, which encodes the multidrug resistance-associated protein MPRP4, known to play a role in paracetamol efflux was also upregulated at Day 20^99^. In another model assessing chronic paracetamol exposure effects in E15-19 rat brains^100^, similar to our findings, an increased expression of genes related to proliferation (e.g. *MKI67, MTDH*), cytoskeletal structural genes (e.g. *NEFL* at Day 20), metabolic stress alleviation (e.g. *PFKP* at Day 7) or decrease in genes related to differentiation (*SOX12* at Day 7) migration and dendrite orientation (such as *CRMP1* at Day 20) was found. Notably, a significant increase in *APOE* expression at P200 cells at Day 7 was observed indicating stress or neuronal damage^101^, and a similar response by the increased expression for glutamate transporter *EAAT4*^102^. The significant *IL6ST* (GP130) and *MEF2A* ^103^ upregulation and the similarities of our findings to the other in vivo models, validate both the trajectories identified by our approach, and the chosen paracetamol neurotoxicity platform for early development studies.

Abiding by the Findability, Accessibility, Interoperability, and Reusability (FAIR) principles, we provide full access to the scRNA-seq and scATAC-seq datasets in open access shiny web platforms for scRNA-seq (**<u>hescneuroparacet</u>**) and integrative scATAC-seq/scRNA-seq (**<u>hescneurodiffparacet</u>**)^104,105^. These webtools allow for data correlation with other published gene expression datasets, and enable plotting, exporting, and downloading high resolution figures of gene expression and gene co-expression analysis UMAPs of any gene. Furthermore, there are multiple options for dataset exploration and visualization, such as heatmaps, violin-, box-, proportion- and bubble plots in different tabs, where gene expression may be viewed per cell, cluster, treatment and timepoint. Chromatin opening can be explored for genomic loci of interest for input sample, ATAC clusters or integrated clusters in a genome browser view with peak-to-gene link annotations (instantly plotted for 500 random cells each time). In addition, the processed datasets and code are made available for customization and integration with other studies. To our knowledge this is the first in vitro study of paracetamol exposure of hESCs under neuronal differentiation, that provides access to scRNA-seq and scATAC-seq data via interactive webtools.

Our study has several limitations and strengths. Protein expression changes were not explored, as this was beyond the scope of this study. This study aimed to delineate the epigenetic impact of paracetamol on early human cortical developmental events. We used both bulk- and scRNA-seq and comparing these two datasets identified a varying degree of overlapping DEGs from 1-27 %, confirming that both datasets are complementary to detect transcriptomic changes induced by paracetamol. Using a multi-omics approach, we unveiled the diversified transcriptional networks related to paracetamol exposure. We could not accurately classify the maturation properties of the cells, or the proportion of cells in G0 phase, highlighting the necessity of new tools to deconvolute the neuronal G0 phase. The results of scATAC-seq analysis presented here are inherently limited by the relatively low genome-per-cell coverage. That means that some open chromatin regions that could have proved relevant for the individual cells or cell populations, may have been missed. Finally, the in vitro results presented here need to be validated by in vivo models and targeted human dataset exploration in future studies. This could confirm whether the pathways identified can explain the paracetamol-induced adverse effects in the early human brain.

Using an in vitro hESC model of neuronal development, we identified altered gene expression, DNAm and chromatin openness involved in key neuronal differentiation processes at bulk and single-cell RNA levels. An overlap of genes identified in this model and in the cord blood of neonates exposed long-term to paracetamol during pregnancy, points towards altered epigenetic regulation of early brain development. Identification of common DNAm modification sites and chromatin openness regions with a regulatory role on gene expression can identify loci and underlying mechanisms involved in neurotoxicity of drugs on the developing fetus with potential effect on long-term outcomes. Such findings could strengthen causal inference and clinical translation of altered DNAm, such as in cord blood, on brain development. However, these in vitro results need further validation for potential clinical translation.

## Supplemental information

Supplemental Information includes seven figures and four tables.

## Supporting information

SpildrejordeSupplementalInfo

## Acknowledgements

We thank Knut Waagan (University of Oslo) for technical assistance. The majority of informatic analysis was performed at Saga super computing resources (Project NN9632K) provided by UNINETT Sigma2—the National Infrastructure for High Performance Computing and Data Storage in Norway. The sequencing service was provided by the Norwegian Sequencing Centre (www.sequencing.uio.no), a national technology platform hosted by Oslo University Hospital and the University of Oslo and supported by the “Infrastructure” programs of the Research Council of Norway and the Southeastern Regional Health Authorities. PharmaTox Strategic Research Initiative was supported by Faculty of Mathematics and Natural Sciences, University of Oslo. We thank the Norwegian Centre for Stem Cell Research for discussions at the initiation of this project. We would also like to thank Akshay Akshay (University of Bern) for his help and fruitful discussion on single-cell ATAC-seq data analysis and visualization. We acknowledge funding from Research Council of Norway 262484 (R.E.), 241117 (R.L.), and 287953 (A.S.); the Swedish Research Council 2019-01157 (A.S.) and grants from the Swedish Brain FO2019-0087 (A.S.) and the Freemasons Children’s House of Stockholm (A.S.) and BiomatDB+ (Horizon Europe #101058779; AS). We thank University of Oslo for open access publication support. The funders played no role in study design, data collection, data analyses, interpretation, or writing of the article. The graphical abstract was generated in Biorender.com.

## Author Contributions

Conceptualization, M.S., A.S., R.E., K.G. and R.L.; Methodology, A.S., M.S., R.L., S.M., K.G., R.E., A.Sh., M.L., and M.F.; Writing – Original Draft, A.S., M.S., A.Sh., R.E., K.G. and R.L.; Writing – Review & Editing, A.S., M.S. R.L., K.G., R.E., A.Sh., M.L., H.N., and G.A.; Software, Formal Analysis, and Visualization, M.S., A.Sh., K.G., A.Y.M.S., R.L., and R.E.; Investigation and Validation, A.S., M.S., M.L., M.F., S.M., R.E. and K.G.; Funding, R.L., K.G., A.S., and R.E.; Supervision, R.L., K.G., H.N., A.S., and R.E.; Resources G.A., R.L., and R.E. All authors read and approved the final manuscript.

## Declaration of Interest

The authors declare no competing interest.

## Authors’ information

Ankush Sharma is now an employee of Acerta-Pharma, The Netherlands.

## STAR Methods

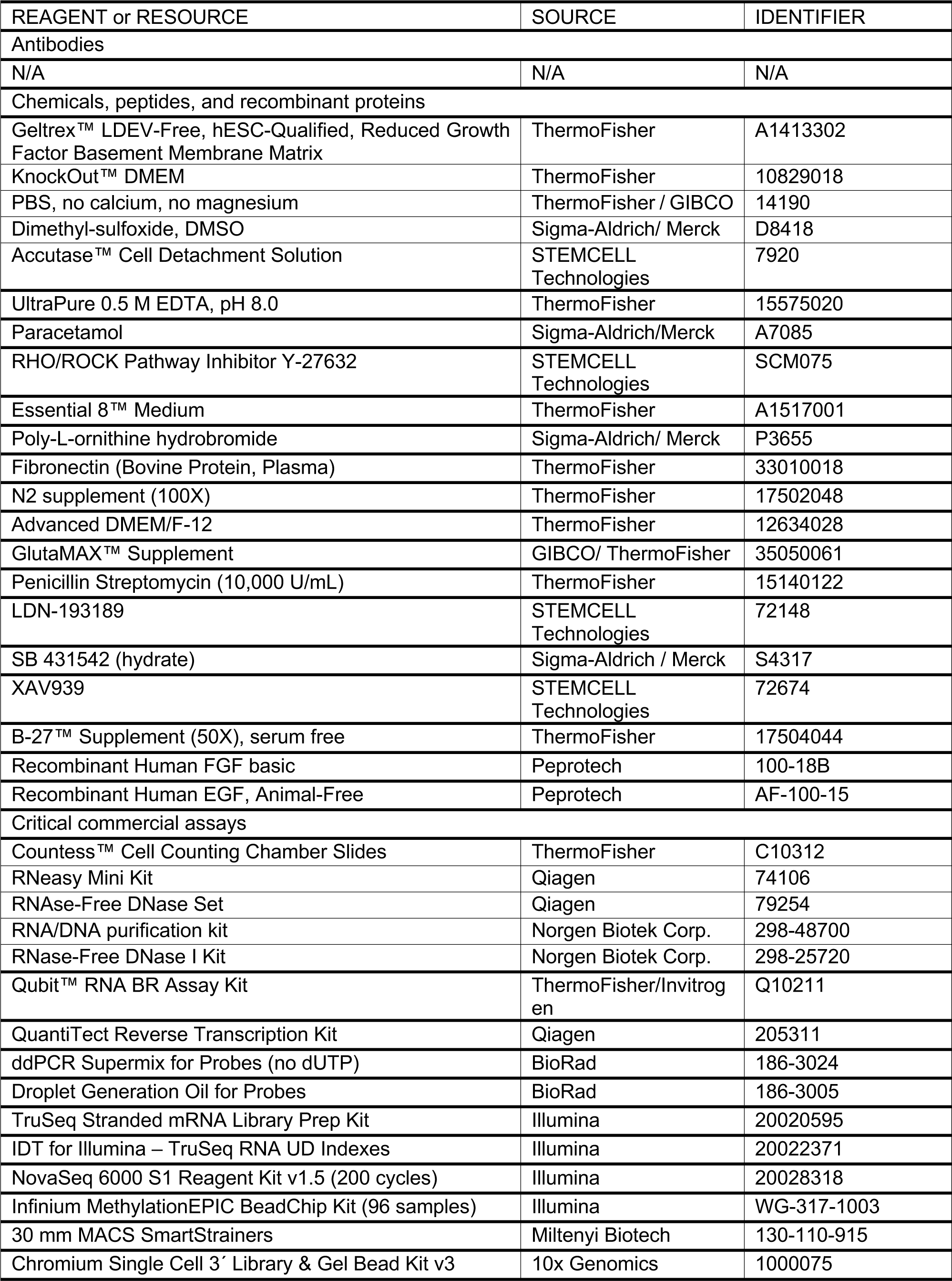

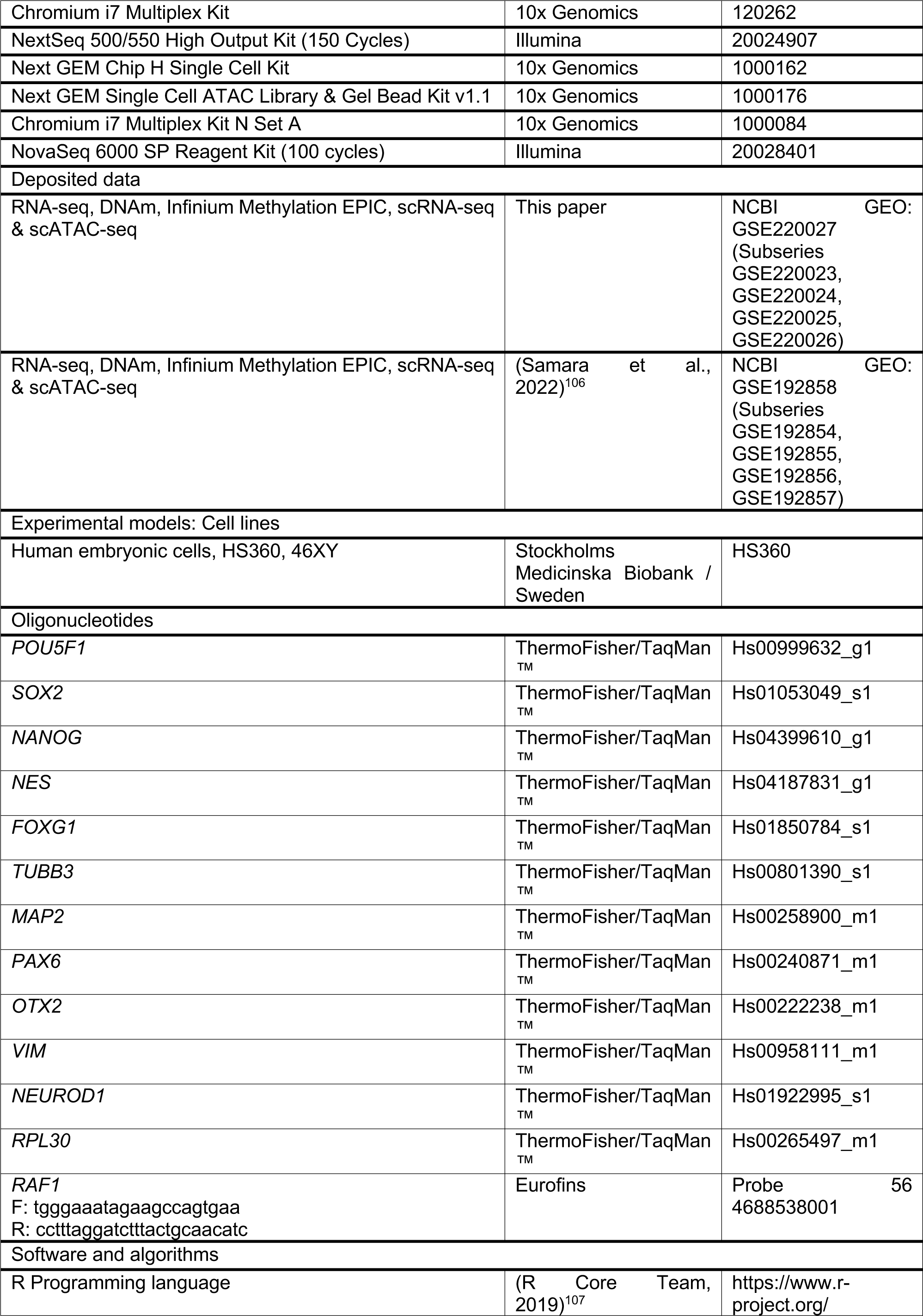

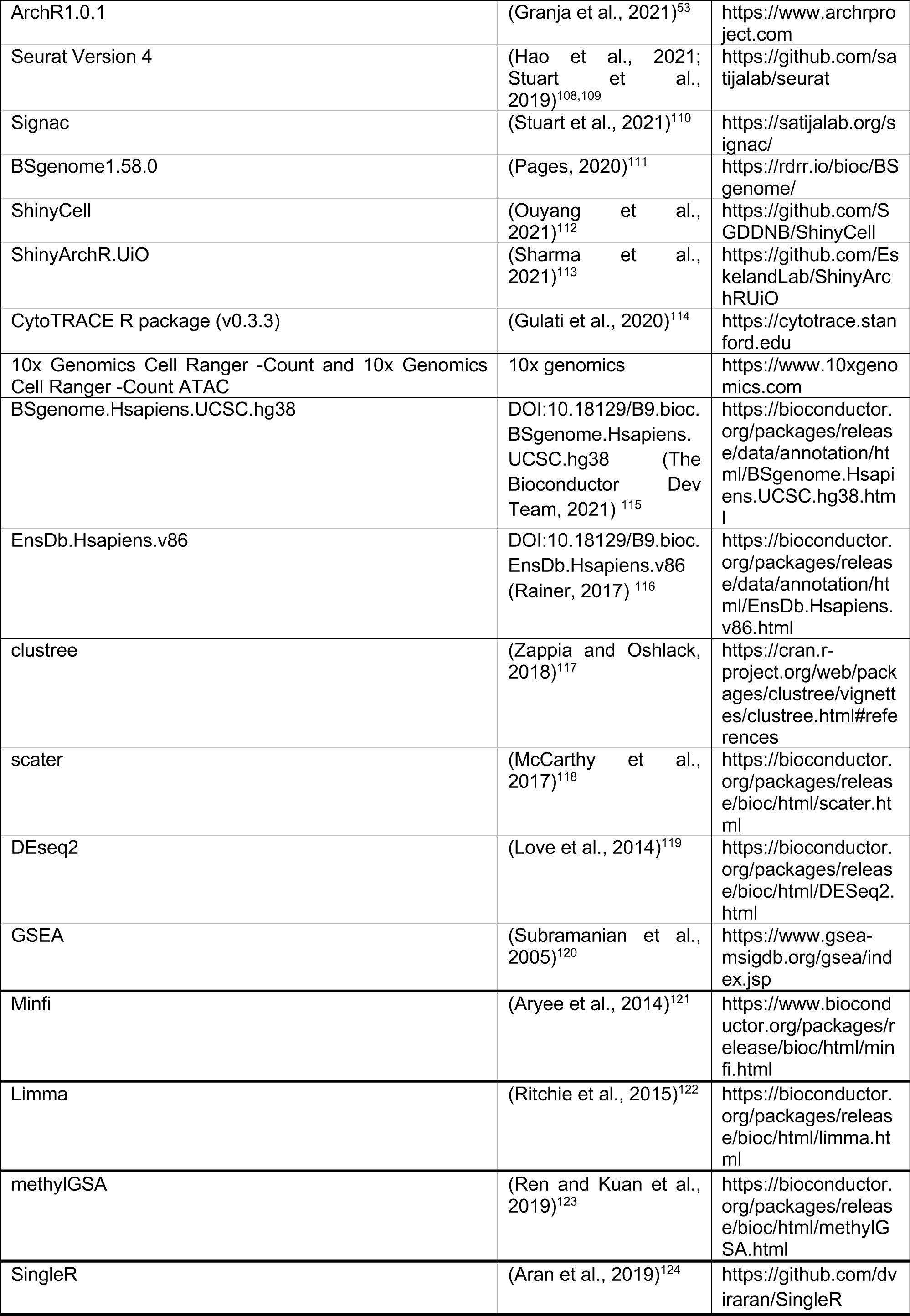

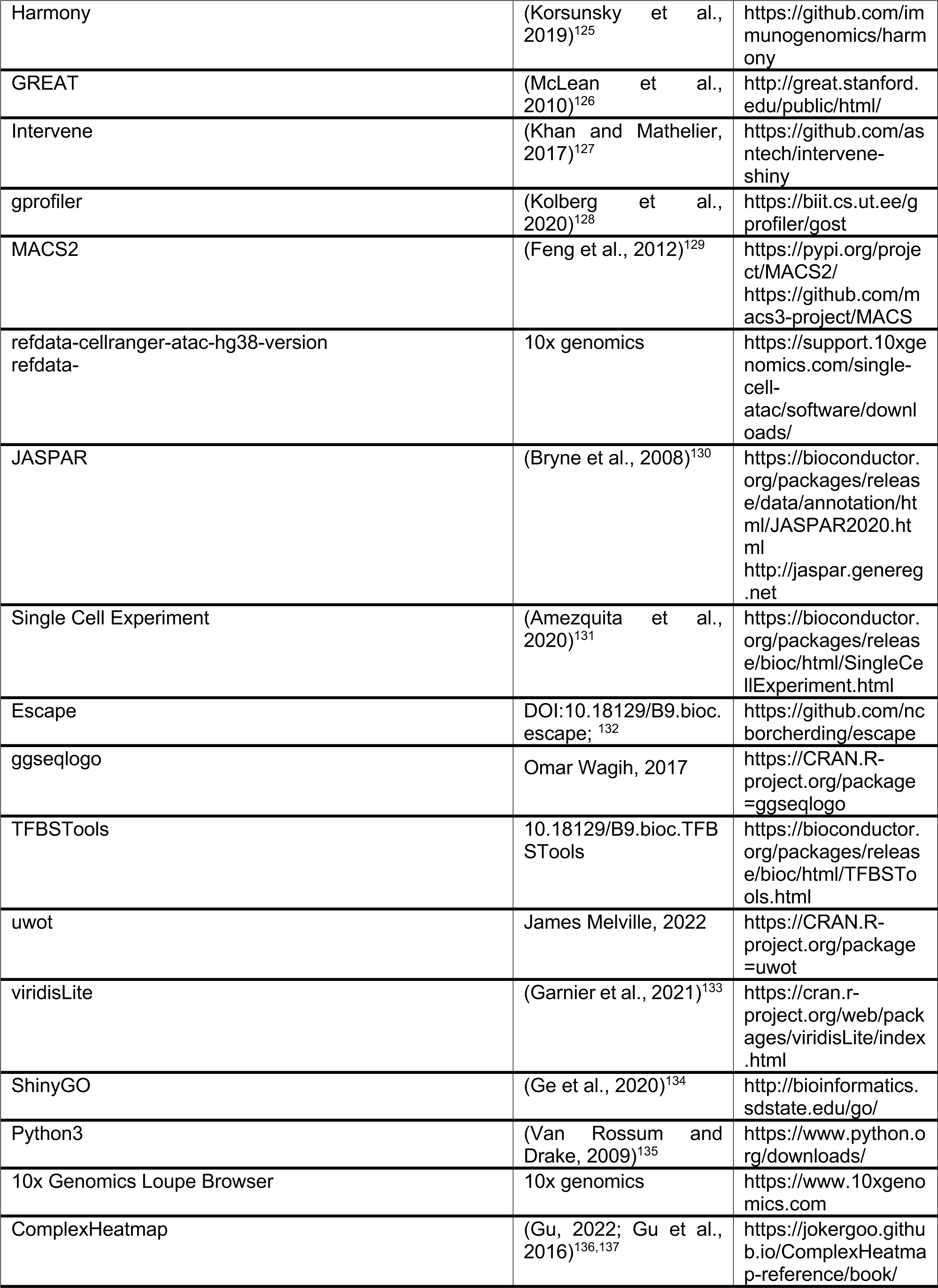

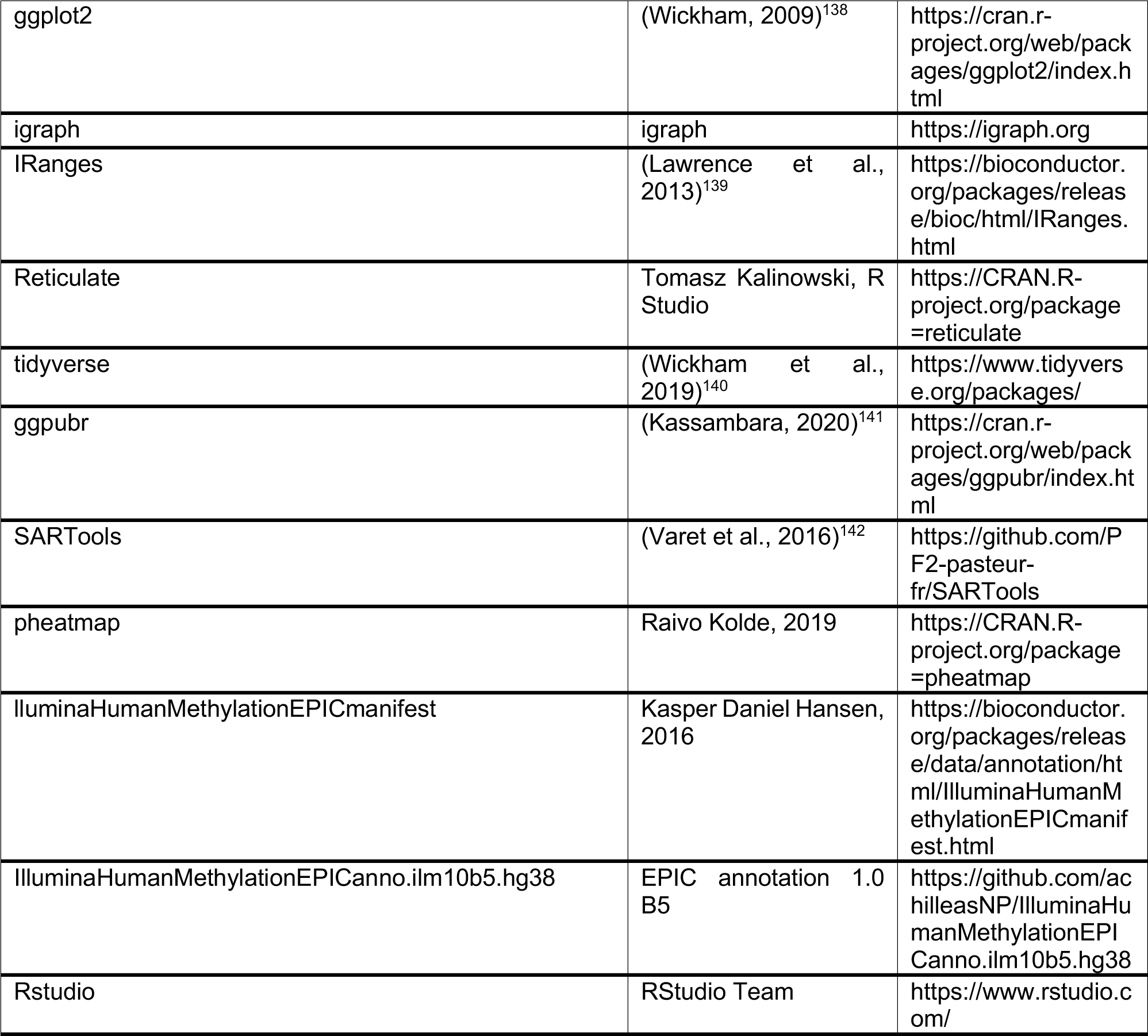

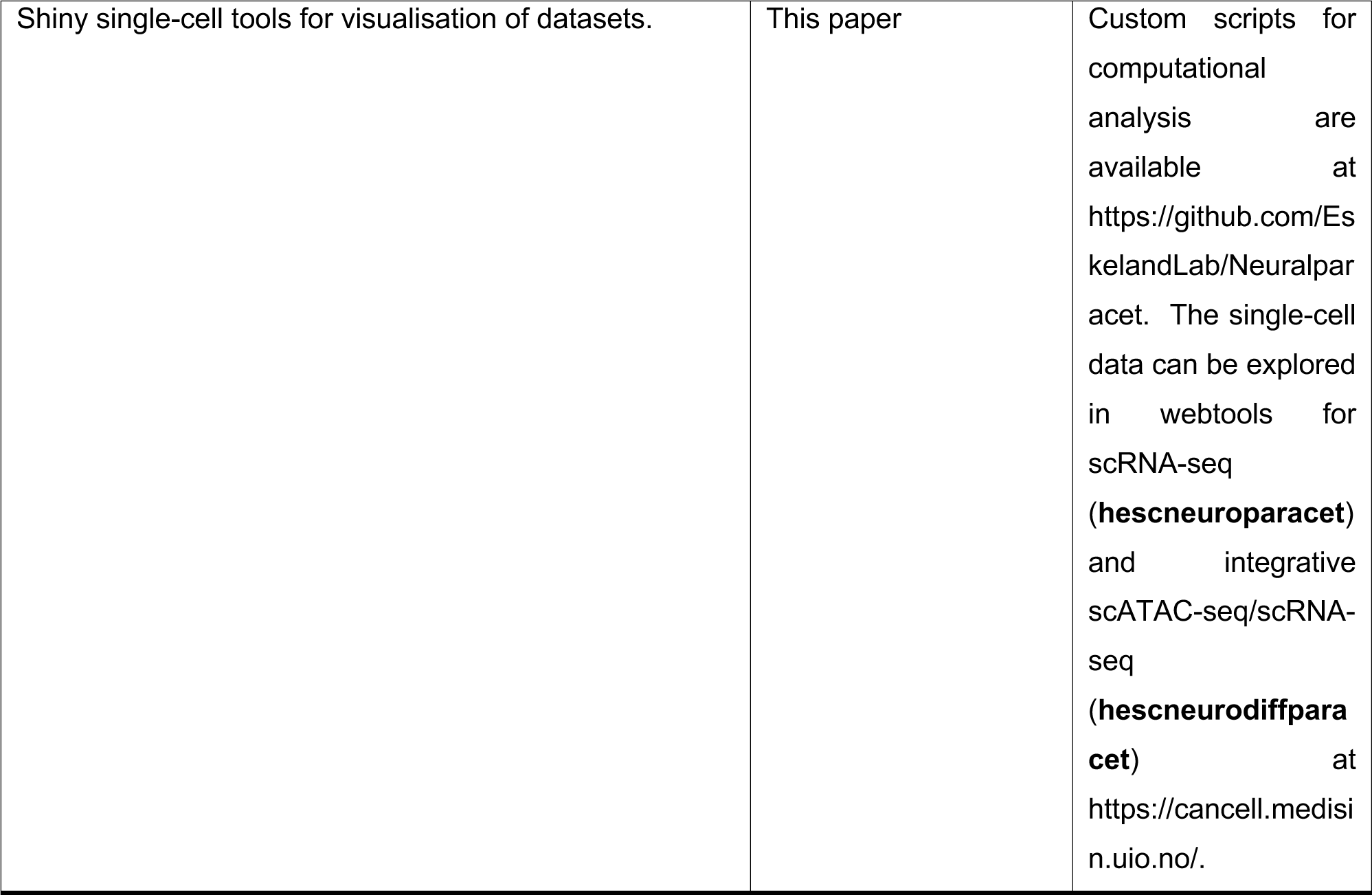

### RESOURCE AVAILABILITY

#### Lead contact

Further information and requests for resources and reagents should be directed to and will be fulfilled by the lead contact Ragnhild Eskeland.

#### Materials availability

This study did not generate new unique reagents.

#### Data and code availability

- All original code can be found at https://github.com/EskelandLab/Neuralparacet. DOIs are listed in the key resources table.
- The single-cell RNA-seq/ATAC-seq, RNA-seq and DNA-methylation data reported in this study cannot be deposited in a public repository because the data could be potentially traced back to a single embryo and the donor. To request access, contact the lead author and the Stockholm Medical Biobank. It may be required to establish a Personal Data processing (PDP) Agreement and/or Data Transfer Agreement (DTA) according to General Data Protection Regulation (GDPR).
- Processed datasets have been deposited at NCBI’s GEO. Accession numbers are listed in the key resources table. Single-cell data are shared for visualization in two open access webtools at https://cancell.medisin.uio.no: (**hescneuroparacet**) https://cancell.medisin.uio.no/scrna/hescneuroparacet/ and (**hescneurodiffparacet**) https://cancell.medisin.uio.no/scatac/hescneurodiffparacet/

### EXPERIMENTAL MODEL AND SUBJECT DETAILS

#### hESC culture and maintenance

hESCs HS360 (Karolinska Institutet, Sweden, RRID:CVCL C202)^86,143^ were cultured as described by Samara et al, 2022^24^. In brief, cells were maintained in Essential 8™ Medium, on Geltrex pre-coated culture plates. Cells were routinely passaged at 75-85% confluency using 0.5 mM ethylenediaminetetraacetic acid (EDTA) in ratios between 1:3 to 1:6. When hESCs were collected for initiation of the differentiation protocol and Accutase was used to detach and dissociate cells.

#### Neuronal differentiation of hESCs and exposure to paracetamol

hESCs HS360 were differentiated, as described^24^, in two separate time-course experiments, to investigate the in vitro effects of paracetamol exposure in physiologically concentrations relevant to long-term exposure in vivo. Since paracetamol crosses the placenta and the blood-brain barrier and has an insignificant plasma protein binding, maternal plasma/serum and cord blood concentrations can be used as estimates for the amount of paracetamol that reaches the developing fetal brain^144–147^. Concentrations of 100 μM and 200 μM paracetamol were chosen as range for physiological plasma peak concentration in vivo, corresponding to one intermediate and one high peak plasma concentration^25–27^. Thus, cells undergoing neuronal differentiation were exposed to changes of medium only (control) or medium containing 100 or 200 μM paracetamol from Day 1 to Day 20. Control cells were harvested at day 0, 7, 13 and 20, while cells exposed to 100 or 200 μM paracetamol were harvested at Day 7 and 20.

### METHODS DETAILS

#### DNA/RNA isolation

Genomic DNA and total RNA were isolated by direct lysis in the culture vessels followed by column-based isolation using RNA/DNA purification kit (Norgen Biotek). RNase-Free DNase I Kit (Norgen Biotek) was applied for on-column removal of genomic DNA contamination from RNA isolates. Five RNA isolates were processed using the RNeasy Mini Kit (Qiagen) followed by DNase-treatment using the RNAse-Free DNase Set (Qiagen). All isolations were performed according to the manufacturer’s instructions. Nucleic acid quantification was performed using Qubit (ThermoFisher Scientific), purity was measured using Nanodrop 2000 (ThermoFisher Scientific), while RNA and DNA integrity was assessed using 2100 Bioanalyzer (Agilent Technologies) and 4200 TapeStation (Agilent Technologies), respectively.

#### ddPCR and RNA expression analysis

Reverse transcription of total RNA was performed using QuantiTect Reverse Transcription Kit (Qiagen). Subsequent ddPCR reactions were set up using ddPCR Supermix for Probes (No dUTP) (BioRad) and Taqman assays (ThermoFisher) or Universal Probes (Roche) in combination with target primers (Eurofins) as outlined in key resources table. Droplets for ddPCR amplification were generated using the QX200 Droplet Generator (BioRad). Data acquisition and primary analysis was done using the QX200 Droplet Reader (BioRad) and QuantaSoft software (BioRad). All steps were performed according to the manufacturer’s instructions. To calculate the number of target copies per ng RNA input, samples were normalized using RPL30 and RAF1 as normalization genes^148^. Statistical comparisons were performed in R using t-test in ggpubr package v.0.4.0^141^. Results were visualized in R using the tidyverse package^140^.

#### Bulk RNA-seq

The sequencing library was prepared with TruSeq Stranded mRNA Library Prep (Illumina) according to manufacturer’s instructions. The libraries (n=39) were pooled at equimolar concentrations and sequenced on an Illumina NovaSeq 6000 S1 flow cell (Illumina) with 100 bp paired end reads. The quality of sequencing reads was assessed using BBMap v.34.56^149^ and adapter sequences and low-quality reads were removed. The sequencing reads were then mapped to the GRCh38.p5 index (release 83) using HISAT2 v.2.1.0^150^. Mapped paired end reads were counted to protein coding genes using featureCounts v.1.4.6-p1^151^. Differential expression analysis was conducted in R version 3.5.1^152^ using SARTools v.1.6.8^142^ and the DESeq2 v.1.22.1 ^153^, and genes were considered significantly differentially expressed with an FDR < 0.05. Normalized counts were visualized using the tidyverse package v.1.3.^140^. The heatmaps were generated using the pheatmap package version 1.0.12^154^. The Wald-test was used to calculate p-values and Benjamini-Hochberg was used to correct for multiple testing. The GSEA analysis of ranked lists of differentially expressed genes were performed using GSEA software v.4.1.0^120^ to identify enrichment of BP terms.

#### Bulk DNAm analysis

DNAm status of 43 samples were assessed using the Infinium MethylationEPIC BeadChip v.1.0_B3 (Illumina). Quality control and pre-processing of the raw data was performed in R using Minfi v.1.36.0^121^. No samples were removed due to poor quality (detection p values >0.05). Background correction was performed using NOOB method ^155^ and β values (ratio of methylated signal divided by the sum of the methylated and unmethylated signal) were normalized using functional normalization ^156^. Probes with unreliable measurements (detection p values >0.01) (n = 12,538) and cross-reactive probes (45) (n = 42,844) were then removed, resulting in a final dataset consisting of 810,477 probes and 43 samples. Probes were annotated with Illumina Human Methylation EPIC annotation 1.0 B5 (hg38). Differential DNAm analysis was performed on the M values (log2 of the β values) using the limma package^122^, and CpGs were considered significantly differentially methylated with an FDR < 0.05. GO analysis of BP terms was performed using p-values of increasing or decreasing CpGs (DMCs) as input to methylRRA function implemented in the methylGSA package version 1.14.0^123^.

#### Collection of cells and scRNA-seq

Cells were washed twice in wells with 1x PBS and detached using Accutase (STEMCELL Technologies) at 37 °C for 7 min. Cells were triturated 10-15 times to separate into single cells and transferred to centrifuge tubes containing the appropriate base media with 0.05 % BSA (Sigma-Aldrich). Counts were performed using Countess II FL Cell Counter (ThermoFisher Scientific) before cells were centrifuged at 300x g for 5 min and the supernatant was discarded. Cell pellets were then resuspended in base medium containing 0.05 % BSA and cell aggregates were filtered out using MACS SmartStrainers (Miltenyi). The cells were recounted and processed within 1 hour on the 10x Chromium controller (10x Genomics). Approximately 2,300 cells were loaded per channel on the Chromium Chip B (10x Genomics) to give an estimated recovery of 1,400 cells. The Chromium Single Cell 3’ Library & Gel Bead Kit v3 (10x Genomics) and Chromium i7 Multiplex Kit (10x Genomics) were used to generate scRNA-seq libraries, according to the manufacturer’s instructions. Libraries from 16 samples were pooled together based on molarity and sequenced on a NextSeq 550 (Illumina) with 28 cycles for read 1, 8 cycles for the I7 index and 91 cycles for read 2. For the second sequencing run, libraries were pooled again based on the number of recovered cells to give a similar number of reads per cell for each sample (33,000 - 44,000 reads/cell).

#### scRNA-seq data analysis

The Cell Ranger 3.1.0 Gene Expression pipeline (10x Genomics) was used to demultiplex the raw base-call files and convert them into FASTQ files. The FASTQ files were aligned to the GRCh38 human reference genome, and the Cell Ranger Count command quantified single-cell read counts using default parameters for Days 7 and 20. Cell ranger aggregate was used for aggregating counts of replicates. The Seurat Package v.4.1.0^109^ was used to perform quality control and normalization on the count matrices. We filtered cells with number of RNA counts >= 2000 & <= 50000 to remove dead cells, doublets and multiplets. The cells expressing fewer genes (less than 200) and genes expressed in less than 3 cells were excluded from the downstream analysis. Outlier cells with high Mitochondrial percentage was computed using Scater package 1.0.4 “isoutlier” function with nmads parameter of 5^118^. Counts were adjusted for cell-specific sampling using the SCTransform function with regression of Cell cycle genes and Mitochondrial content^157,158^.

We used a resolution of 0.4 to cluster cells, obtained by determining the optimum number of clusters (cells grouped together sharing similar expression profiles) in the dataset using the Clustree R package^117^. Principal component analysis was performed using the RunPCA function, followed by FindClusters and RunUMAP functions of Seurat package to perform SNN-based UMAP clustering.

FindMarkers from the Seurat R package were used to perform differential expression analysis between groups. For DE between exposure groups, thresholds were set to the following: min.pct = 0.25, min.diff.pct = -Inf, logfc.threshold = 0.1. For top overlapping DE genes per Day (Fig. 6 E/J), thresholds were set to the following: min.pct = 0.25, min.diff.pct = -Inf, logfc.threshold = 0.25. Genes with an adjusted p-value < 0.05 were considered significant. GO analysis was performed using the DEenrichRPlot function of the mixscape R package with the "GO_Biological_Process_2018" database with the following thresholds: logfc.threshold = 0.25, max genes = 500. The SingleR R package ^124^ was used to annotate the cells against reference data sets from a Human Brain dataset ^46^ from the scRNAseq R package ^159^. Cell types with < 15 cells annotated were excluded from the plots.

#### scATAC-seq Library preparation and sequencing

Cells were washed twice with 1xPBS and detached to single cell suspension by application of Accutase (STEMCELL Technologies) at 37 °C for 7 min. The detached cells were washed with appropriate base media with added 0.04% BSA (Sigma-Aldrich) and filtered using MACS SmartStrainers (Miltenyi Biotech) to remove cell aggregates. Nuclei isolation was done according to the 10x Genomics protocol CG000169 (Rev D) using 2 minutes of incubation in a lysis buffer diluted to 0.1x and 0.5x for Day 0 and Day 20 cells, respectively. Countess II FL Cell Counter (ThermoFisher Scientific) was used to quantify nuclei and confirm complete lysis and microscopy to confirm high nuclei quality. Nuclei were further processed on the 10x Chromium controller (10x Genomics) using Next GEM Chip H Single Cell Kit (10x Genomics), Next GEM Single Cell ATAC Library & Gel Bead Kit v1.1 (10 x Genomics) and Chromium i7 Multiplex Kit N Set A (10x Genomics) according to the Next GEM Single Cell ATAC Reagent Kits v1.1 User Guide (CG000209, Rev C). The targeted nuclei recovery was 5,000 nuclei per sample. The resulting 4 sample libraries were sequenced on a NovaSeq Sp flow cell (Illumina) with 50 cycles for read 1, 8 cycles for the i7 index read, 16 cycles for the i5 index read and 49 cycles for read 2.

#### scATAC sequencing analysis

Cell Ranger ATAC version 1.2.0 with reference genome GRCh38-1.2.0 was used to pre-process scATAC-seq raw sequencing data into FASTQ files. Single-cell accessibility counts for the cells were generated from reads using the *cellranger-atac count* pipeline. Reference genome HG38 used for alignment and generation of single-cell accessibility counts was obtained from the 10x Genomics (https://support.10xgenomics.com/single-cell-atac/software/downloads/).

Downstream analysis of the scATAC-seq data was performed using the R package ArchR v1.0.1^53^. A tile matrix of 500-bp bins was constructed after quality control, removal of low-quality cells and doublet removal using the doubletfinder function of ArchR. The ArchR Project contained the filtered cells that had a TSS enrichment below 3 and <1000 fragments. A layered dimensionality reduction approach utilizing Latent Semantic Indexing (LSI) and Singular Value Decomposition (SVD) applied on Genome-wide tile matrix. Uniform Manifold approximation and projection (UMAP) was performed to visualize data in 2D space. Louvain Clustering methods implemented in R package Seurat^109^ were used for clustering of the single-cell accessibility profiles.

### QUANTIFICATION AND STATISTICAL ANALYSIS

Statistical analyses were performed in R version 4.1.2^107^ applying SARTools v.1.6.8^142^, DESeq2 v.1.22.1^119^, Limma^122^, ArchR^53^, Seurat^108,109^, and ggpubr package v.0.4.0 ^141^. Details are described in the relevant methods sections above.

## Notes

https://cancell.medisin.uio.no/scrna/hescneuroparacet/

https://cancell.medisin.uio.no/scatac/hescneurodiffparacet/

### Competing Interest Statement

The authors have declared no competing interest.

### Summary of Updates

Manuscript and Supplemental files updated.

