## Supplementary material for "Multi-omics approach reveals dysregulated genes during hESCs neuronal differentiation exposure to paracetamol": SpildrejordeSupplementalInfo

[1]PharmaTox Strategic Research Initiative, Faculty of Mathematics and Natural Sciences, University of Oslo, Norway
[2]Department of Medical Genetics, Oslo University Hospital and University of Oslo, Norway
[3]Institute of Clinical Medicine, Faculty of Medicine, University of Oslo, Oslo, Norway.
[4]Division of Clinical Paediatrics, Department of Women's and Children's Health, Karolinska Institutet, Sweden
[5]Astrid Lindgren Children's Hospital, Karolinska University Hospital, Stockholm, Sweden
[6]Department of Informatics, University of Oslo, Norway
[7]Department of Molecular Medicine, Institute of Basic Medical Sciences, Faculty of Medicine, University of Oslo, Oslo, Norway
[8]Department of Biosciences, University of Oslo, Norway
[9]Division of Obstetrics and Gynecology, Department of Clinical Science, Intervention and Technology (CLINTEC), Karolinska Institutet, Alfred Nobels Allé 8, SE-14152, Stockholm, Sweden.
[10]Center for Fetal Medicine, Karolinska University Hospital, SE-14186 Stockholm, Sweden.
[11]Pharmacoepidemiology and Drug Safety Research Group, Department of Pharmacy, University of Oslo, Norway
[12]Division of Clinical Neuroscience, Department of Research and Innovation, Oslo University Hospital, Oslo, Norway
[13]Centre for Fertility and Health, Norwegian Institute of Public Health, Oslo, Norway
[14]Lead contact

[#]Current address: Department of Analysis and Diagnostics, Section for Molecular Biology, Norwegian Veterinary Institute, Ås, Norway
[§]Current address: Department of Medical Genetics, Oslo University Hospital and University of Oslo, Norway
[+]Current address: Istituto di Genetica Molecolare, CNR - Consiglio Nazionale delle Ricerche, Pavia, Italy.

### Supplemental Figures

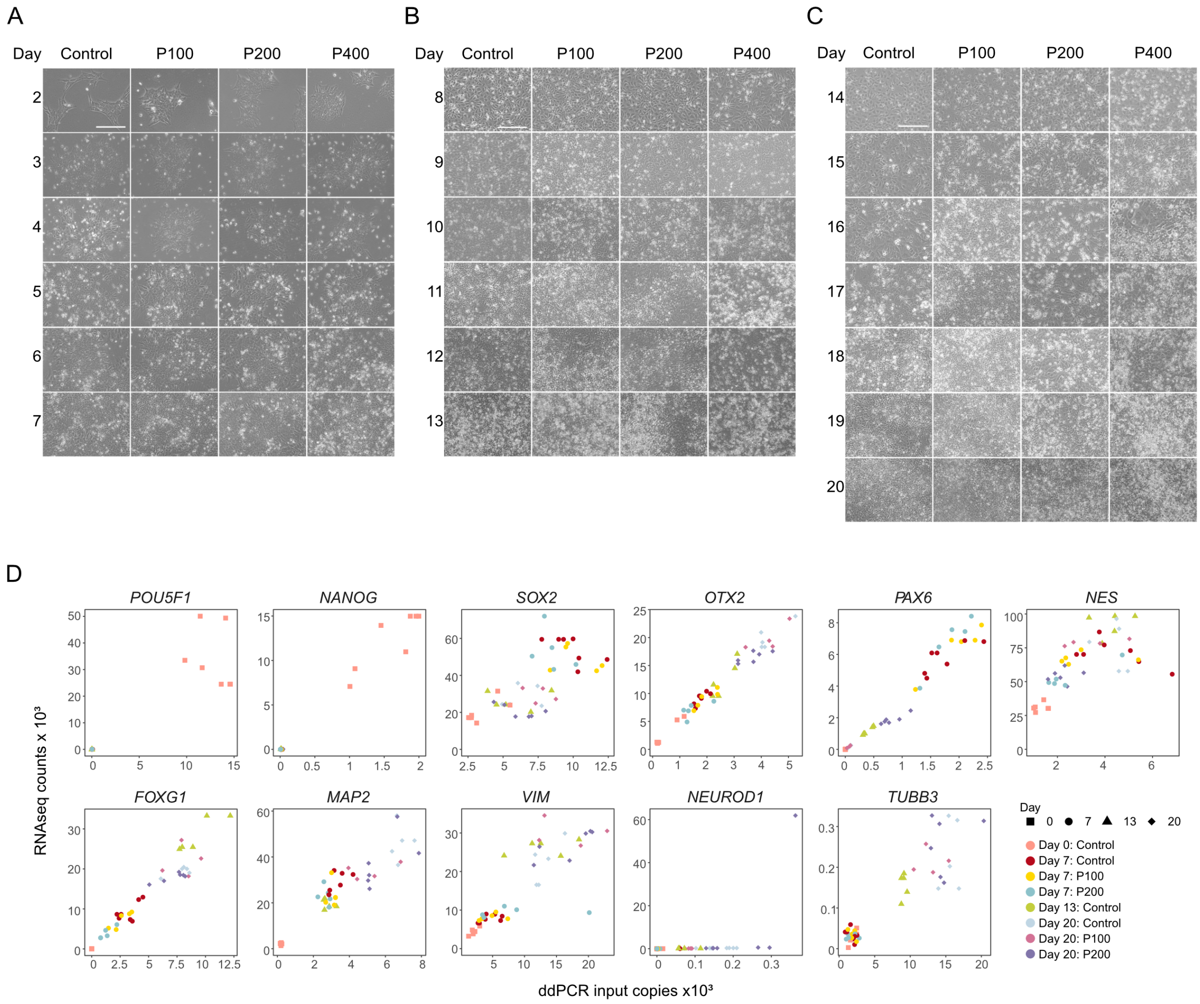


**Figure S1. Differentiation timeline for control cells and cells treated with 100, 200 or 400 µM paracetamol.** Brightfield images of control cells, P100, P200 or 400 µM paracetamol (P400) differentiation A) Day 2-7, B) Day 8-13 and C) Day 14-20. Images were taken with an EVOS FL microscope at 20X magnification (scale bar corresponds to 100 µm). D) ddPCR input copies versus bulk mRNA expression of selected marker genes from Day 0, 7, 13 and 20 in control cells and cells exposed to 100 or 200 µM paracetamol. One dot represents one replicate.

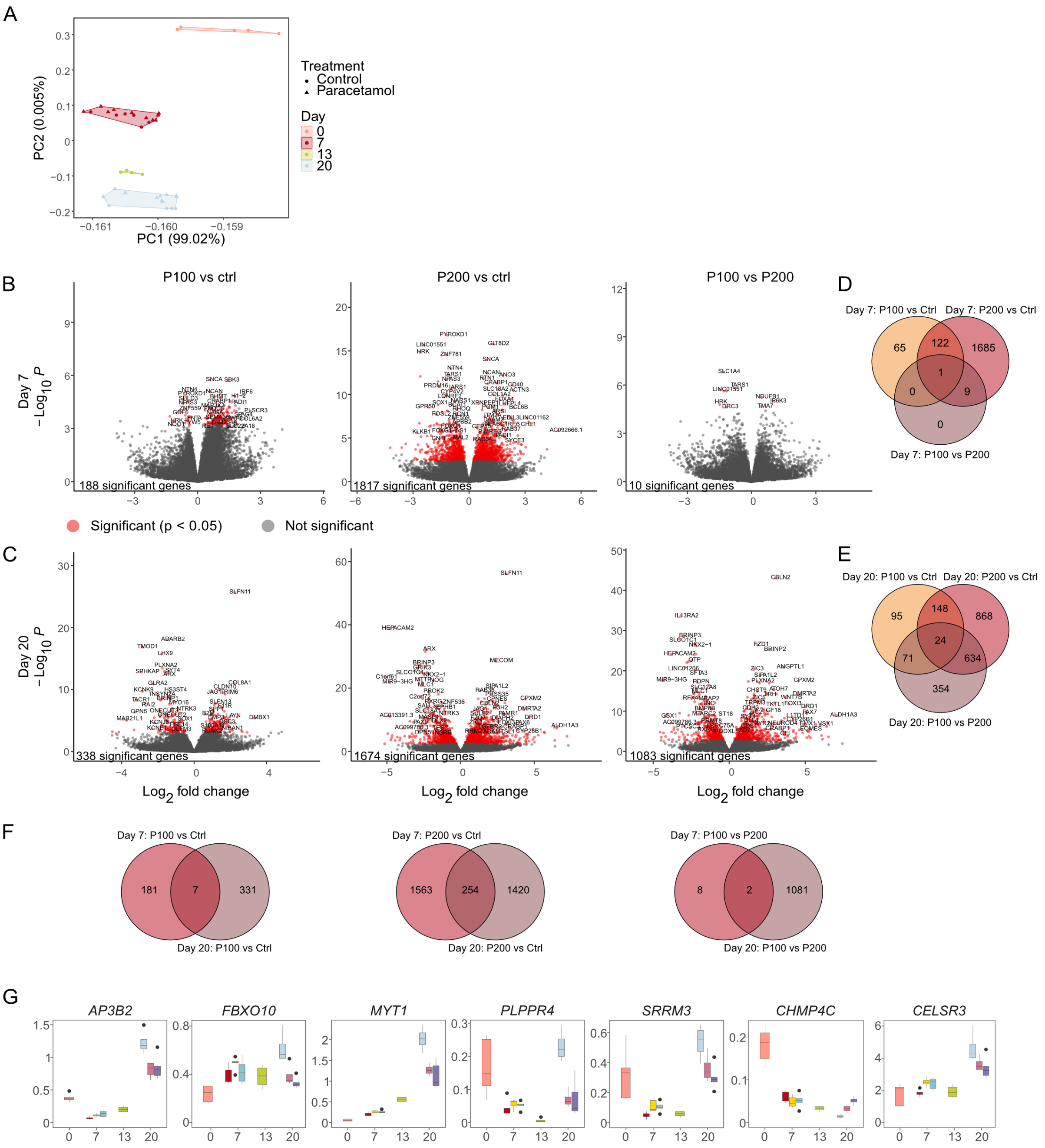


**Figure S2. Global gene expression analysis of differentiating neuronal cells exposed to paracetamol.** A) Principal component analysis of replicates coloured by day and exposure group. B-C) Volcano plots showing differentially expressed genes between cells treated with 100 µM paracetamol compared to control (left), 200 µM paracetamol compared to control (middle) and 100 µM paracetamol compared to 200 µM paracetamol (right) at B) Day 7 and C) Day 20. D-F) Venn diagrams show number of overlapping DMCs between D) Day 7 comparisons, E) Day 20 comparisons and F) Day 7 and Day 20 comparisons. Genes with FDR < 0.05 are considered significant.

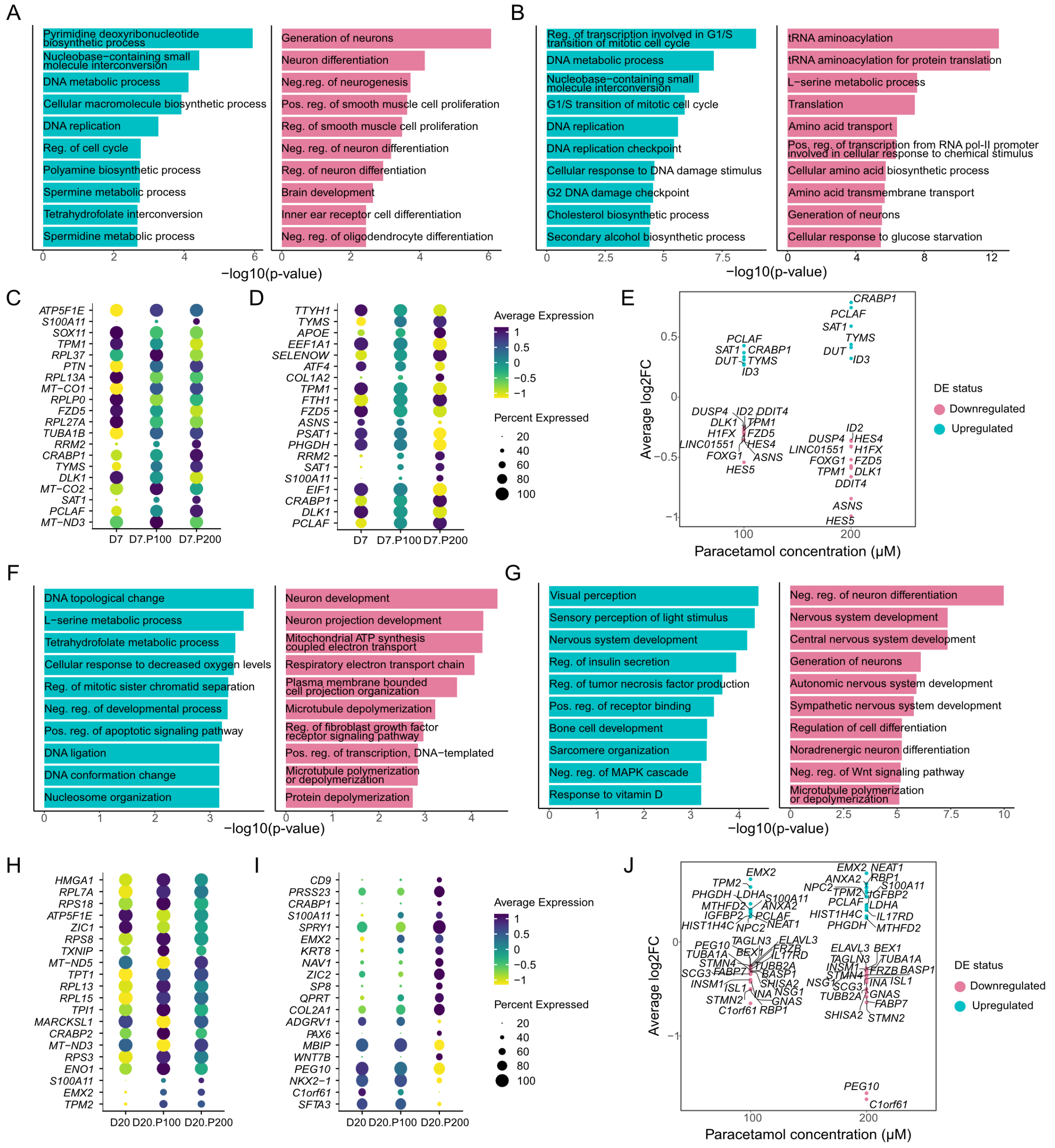


**Figure S3. Differential gene expression analysis in single cells showed downregulation of genes involved in neuronal differentiation after paracetamol exposure.** A-B) Top 10 upregulated (green) and downregulated (pink) BPs among DEGs at Day 7 between A) P100 or B) P200 and control cells. C-D) Bubble plot showing gene expression for the top 20 DEGs at Day 7 between C) P100 or D) P200 and control cells. E) Gene expression of top overlapping genes between P100 and P200 cells compared to control cells at Day 7. F-G) Top 10 upregulated (green) and downregulated (pink) BPs of DEGs at Day 20 between F) P100 or G) P200 and control cells. H-I) Bubble plot showing gene expression for the top 20 DEGs at Day 20 between H) P100 or I) P200 and control cells. J) Gene expression of top overlapping genes between P100 and P200 cells compared to control cells at Day 20.

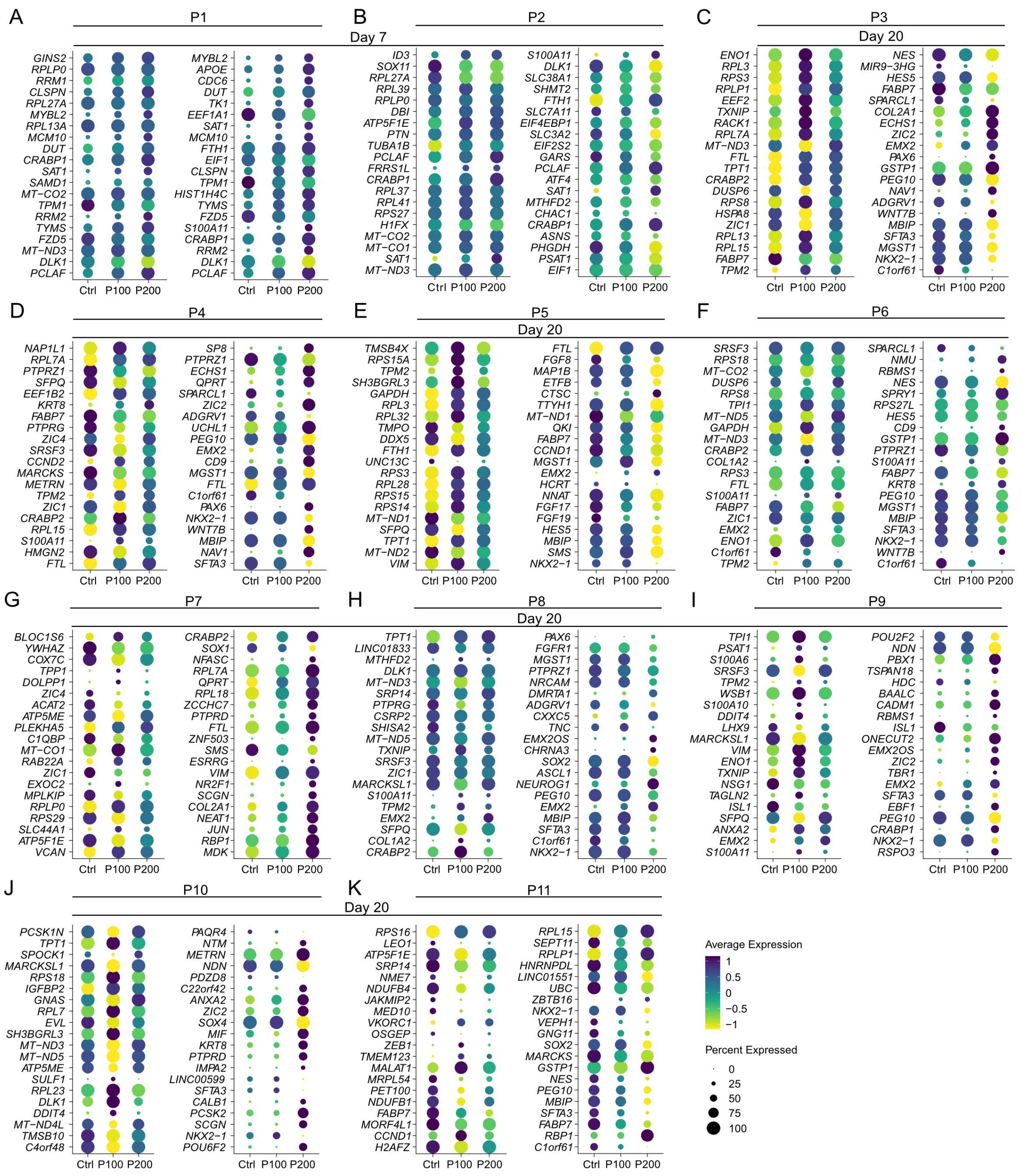


**Figure S4. Single-cell differential expression analysis per cluster.** Gene expression of the top 20 differentially expressed genes between P100 cells and control (left) and P200 cells and control (right) in cluster A) P1, B) P2, C) P3, D) P4, E) P5, F) P6, G) P7, H) P8, I) P9, J) P10 and K) P11.

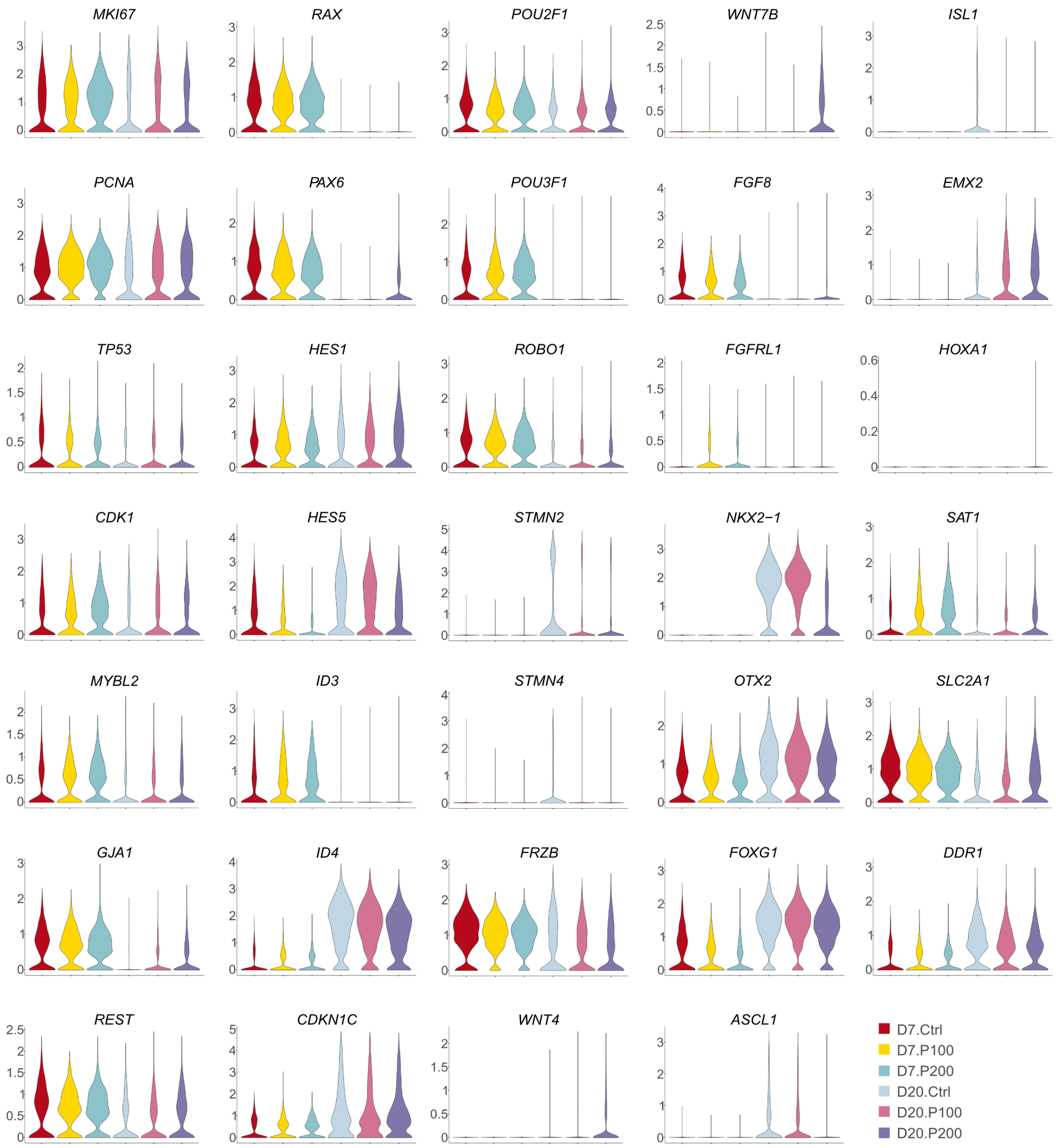


**Figure S5. Gene expression changes in paracetamol compared to control in scRNA data.** Gene expression of selected differentially expressed genes *MKI167*, *PCNA*, *TP53*, *CDK1*, *MYBL2*, *GJA1*, *REST*, *RAX*, *PAX6*, *HES1*, *HES5*, *ID3*, *ID4*, *CDKN1C*, *POU2F1*, *POU3F1*, *ROBO1*, *STMN2*, *STMN4*, *FRZB*, *WNT4*, *WNT7B*, *FGF8*, *FGFRL1*, *NKX2-1*, *OTX2*, *FOXG1*, *ASCL1*, *ISL1*, *EMX2*, *HOXA1*, *SAT1*, *SLC2A1* and *DDR1*.

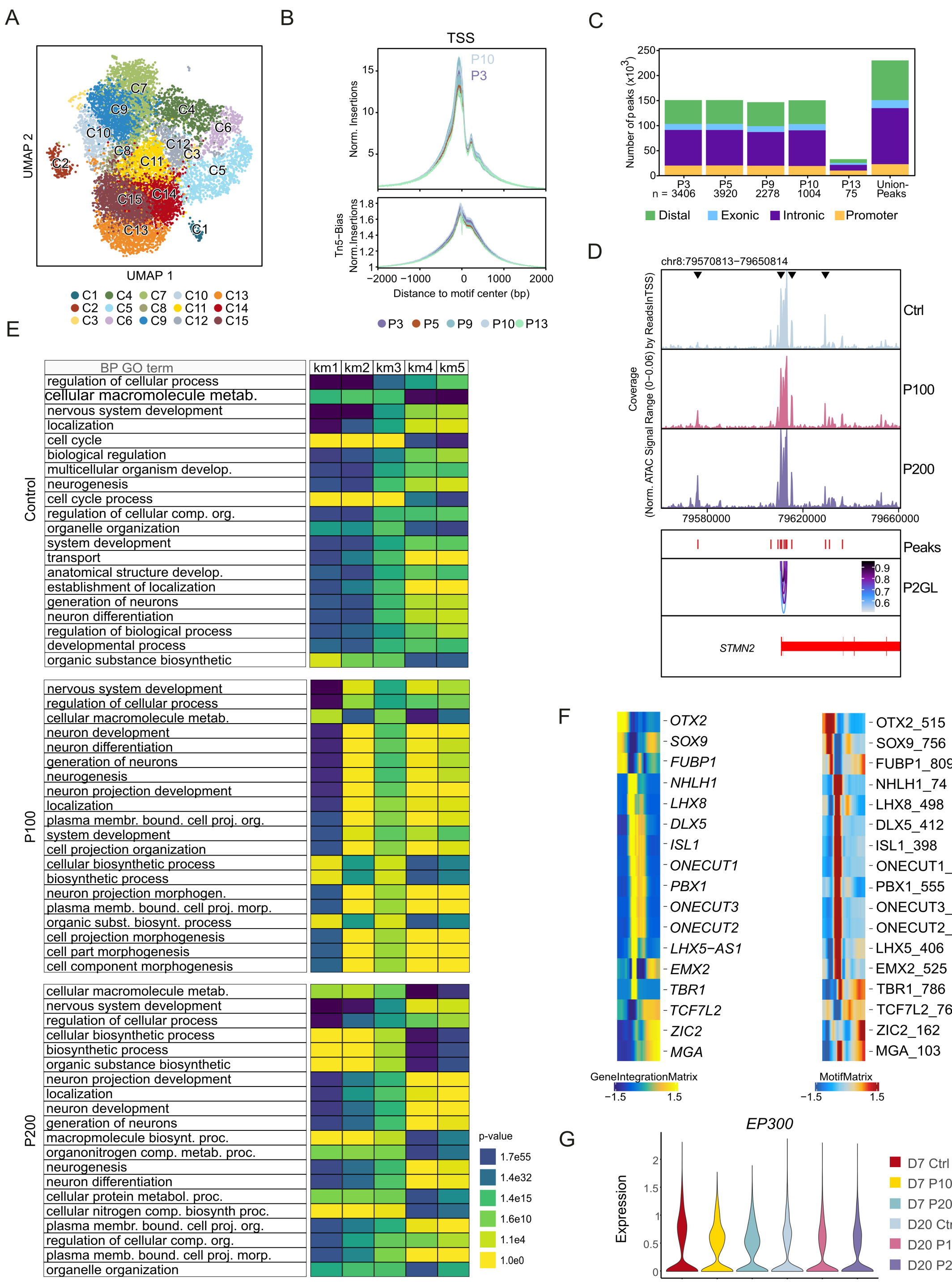


**Figure S6. Integrative chromatin accessibility analysis in neuronal differentiation Day 20 upon treatment with paracetamol.** A) A UMAP plot representing clusters at scATAC-seq modality (C1-C15). B) Chromatin opening across all gene TSS in integrated clusters P3, P5, P9, P10 and P13. Tn5 bias normalized insertions are shown below. C) Bar plot of number of peaks in distal genomic regions, exons, introns and promoters. D) *STMN2* locus browser view (GRCh38.p13) with ATAC-seq signals generated from 5000 cells for control, P100 and P200. ATAC peaks (red) and P2GLs (arcs, blue gradient) are shown. Black arrows indicate some peaks with observed change in ATAC-seq signals between control, P100 and P200. E) Top 20 significant GO terms of controls, P100 and P200 linked genes (genes linked with putative CREs having correlation value greater than 0.45 and with significant FDR<1e-4) for k - mean groups 1-5. F) Top selected motifs for transcription factors in highly active genes shown on a heatmap computed on Geneintegration and Motif Matrix for all Day 20 cells. G) scRNA gene expression of EP300 at Day 7 and Day 20.

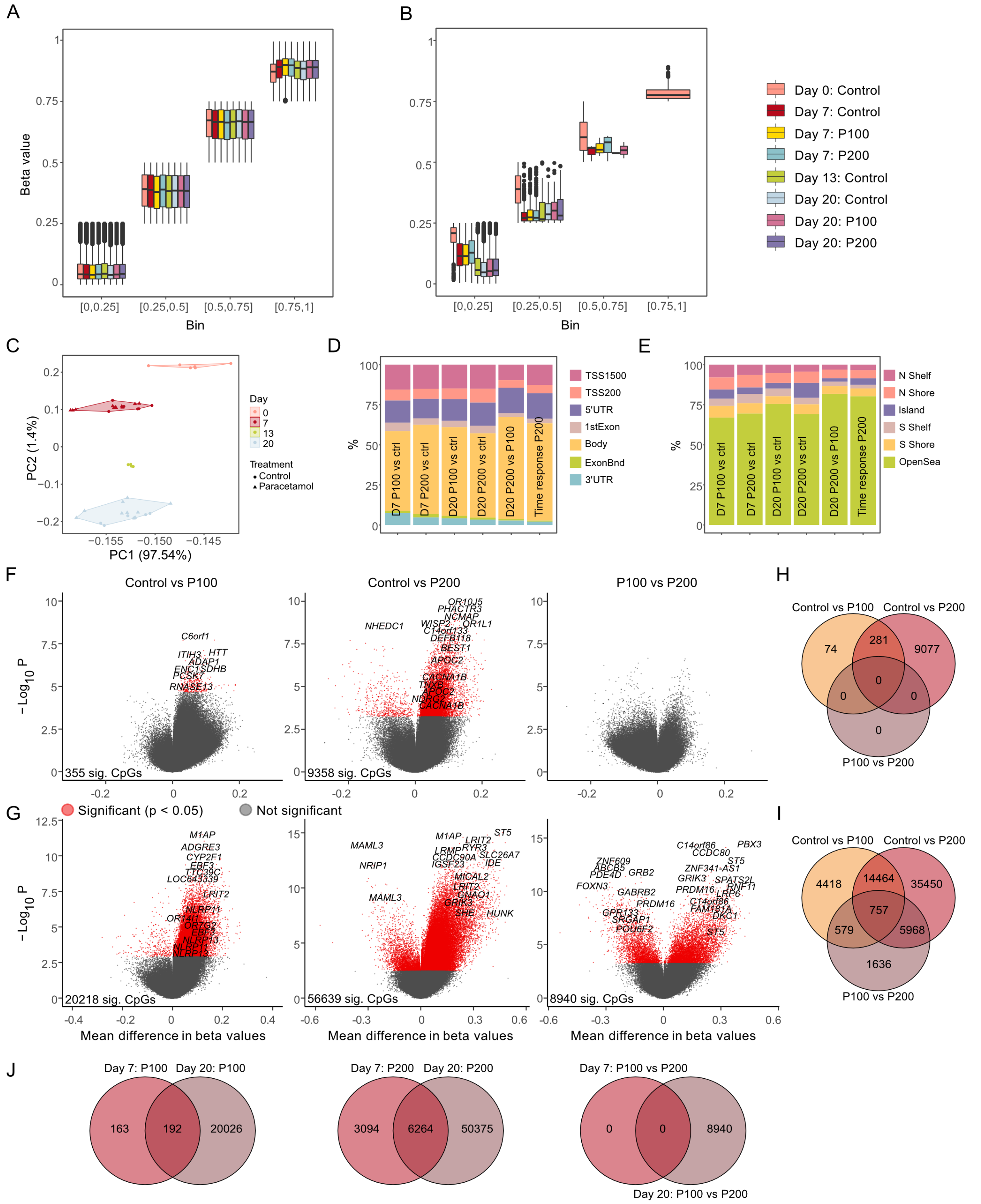

**Figure S7. DNA methylation analysis of neuronal differentiation during paracetamol exposure.** A) Average DNAm levels for each sample across all CpGs and non-CpGs (grouped in bins of 0.25) at different days in controls and cells exposed to different paracetamol doses. B) Average DNAm levels for each sample across all non-CpGs (grouped in bins of 0.25) for all controls and exposed cells.C) Principal component analysis of replicates coloured by day and exposure group. Distribution of significant CpGs in relation to D) annotated genes and E) CpG islands for the different comparisons. F-G) Volcano plots showing DMCs between P100 cells compared to control (left), P200 compared to control (middle) and P100 compared P200 (right) at D) Day 7 and E) Day 20. CpGs with FDR > 0.05 are considered significant. H-J) Venn diagrams show number of overlapping DMCs between H) Day 7 comparisons, I) Day 20 comparisons and J) Day 7 and Day 20 comparisons. CpGs with FDR < 0.05 are considered significant.
